# Membrane voltage multistability in coupled glial cells

**DOI:** 10.64898/2026.05.03.722503

**Authors:** Predrag Janjic, Dimitar Solev, Min Zhou, Ljupco Kocarev

## Abstract

Growing interest in describing the electrical behavior of glial cells, mainly astrocytes in the brain, poses challenges to the commonly accepted belief that they electrically respond only in a linear manner when taking up excess extracellular potassium and glutamate, thereby maintaining the isopotentiality and glutamate level of the networks. Their highly conductive mutual interconnections via gap junction (GJ) connections introduce yet another class of nonlinear elements. As more studies report nonlinearities in both membrane and junctional conductances, the need to formulate minimal dynamical models of their transient behavior is getting more acute. Because ODE models of coupled cells require a simplified description to keep the parameter set small, the still-rare quantitative studies on astrocytes make the task even harder. This study attempts to qualify a self-coupled cell, or a glial cell coupled to a fixed voltage, as a useful system for detecting the nature of membrane voltage *V* instabilities and transitions arising from coupling. In a novel biophysical model of a coupled astrocyte, we introduce nonlinear kinetics of inactivation of the coupling for large junctional voltages, for the first time. We found that N-shaped nonlinearities and the corresponding bistability in isolated-cell dynamics serve as a baseline, on top of which nonlinear coupling enriches the bifurcation picture. Numerical simulations of a 1-D array of coupled astrocytes show that coupling increases the propensity of astrocytic *V* to exhibit bistability and front propagation. We believe the illustrated effects of nonlinear coupling will motivate biologists to further explore their impact on health and disease.

**Significance statement:** Transient changes in membrane voltage of glial cells may produce significant transient voltage differences between directly coupled cells. The nonlinear steady-state conductance of their interconnection elements, gap junctions, introduces nonlinear current profiles that are difficult to measure and quantify with available methods because of the junctions’ high permeability and the general leakiness of glial membranes. We propose a minimal model of the glial membrane extended with a self-coupled feedback loop, which, under realistic simplifying assumptions, could serve to analyze the impact of coupling on the stability of resting membrane voltage qualitatively.

**Author’s summary:** Neuronal cells in the brain and spinal cord cannot exist or function without the supportive and neuromodulatory functions of the diverse glial cell population. This applies to virtually all physiological processes at the cellular level, from cell development, metabolic support, membrane signaling, slow molecular signal transduction, ion homeostasis, and neurovascular coupling to myelination, to name only a few. They all involve neuro-glial interaction. Although all glial cell types are interconnected, the most abundant ones, astrocytes, are extensively interconnected by gap junctions to form ordered networks. Electrically, astrocytic networks display membrane voltage isopotentiality, which is considered the system-wide resting state for a given neuro-glial circuit or unit. With the molecular and cellular substrates of glial connectivity being slowly elucidated, network science and dynamical modeling are slowly “invading” that area, with many important issues left open. In this study, using classical dynamical systems approaches, we indicate how nonlinear intercellular coupling between astrocytes affects the physiological resting state and its instabilities compared with an isolated, uncoupled cell. We hypothesize that this minimal model could fill a gap in ODE modeling of neuro-glial circuits within the broader scope of hypothesis-driven research in cell-level neuroscience.

## 1. Introduction

Glial cells, especially astrocytes, are a diverse class of neural cells that play many roles in neural function, directly interacting with neurons and blood vessels^1,2^ in many brain areas. The crucial systemic feature of astrocytes is their massive interconnection through their extensively branching processes into highly conductive networks, forming a cellular continuum called a *syncytium*^3,4^. Figure 1A shows visualizations of the specific structural properties of astrocytes that largely define their function. Astrocytes form massively interconnected networks^3^ (Fig. 1A and 1B), with each cell directly connecting on average 7–10 immediate neighbors^5^ by electrical junctions formed by gap junction channels (GJCs), located close to the tips of their branches^6^. Each gap junction (GJ) is a cluster of hundreds to thousands of GJCs^7^ (see Supplementary Material S1). The physical size and channel density of GJs^6^ suggest a level of complexity that is difficult to model as a single channel pore^8^.

**Figure 1.**
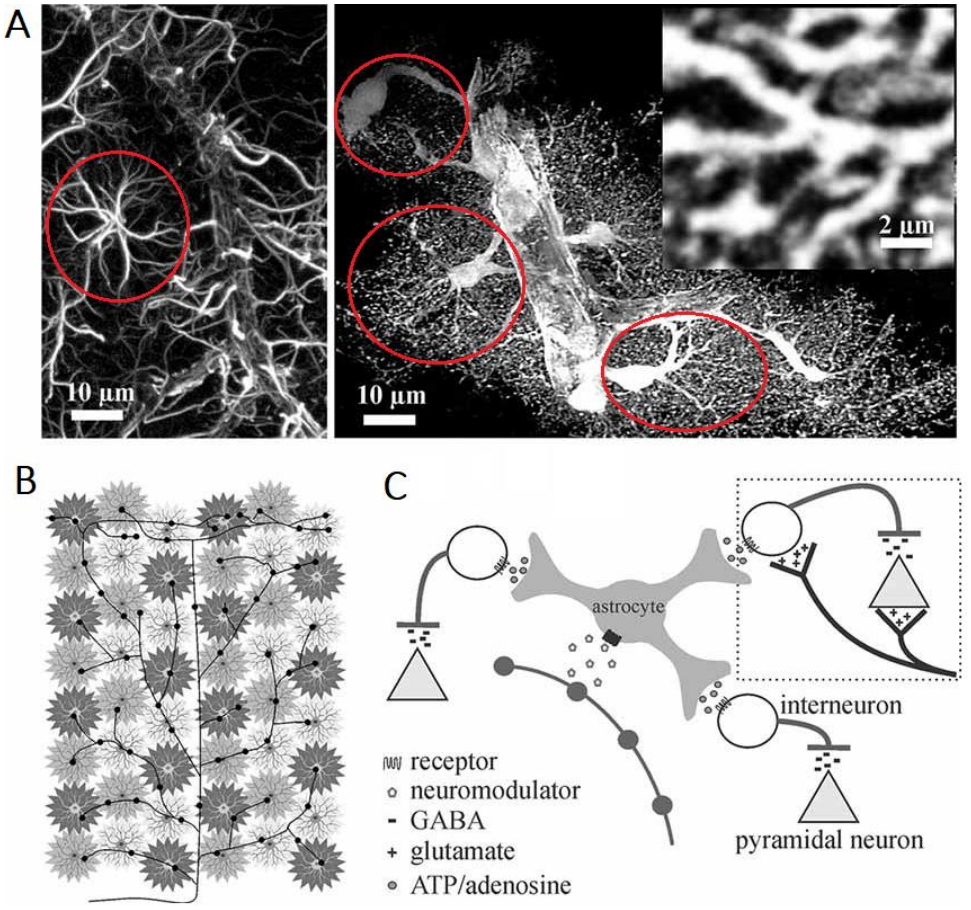
Network features of astrocytes. **(A)** Star-like morphology of astrocytes (circled in red) from the rat cortex, where traditional staining outlines only major branches (left). In contrast, enhanced fluorescence imaging reveals the highly complex and delicate morphology of astrocytes (right). Using their endfeet structure, they connect to the blood vessels (the thick stalks in the right panel in A). Source images adapted from Simard and Nedergaard (2004)^9^, with permission. **(B)** Network patterns of interconnected astrocytes (gray stars) that interact directly with neuronal circuits. **(C)** Neuro-modulatory roles of an astrocyte within a local canonical excitation– inhibition control circuit. Figure adapted from Pacholko et al. (2020)^10^, with permission.

Similar to neuronal cells, the glial membrane displays a wide range of electrical conductances through a variety of ion channels, receptors, transporters, and junctions^4,11,12^. Most of them produce measurable net currents through the membrane, transiently altering the *V* all the time. Figure 1C shows an astrocyte within a typical canonical excitation–inhibition circuit. With the electrical processes in any cell being under dynamical control of *V*, the dynamical stability of *V*_*r*_, the physiological *V* resting state has a profound effect not only on the electrical phenomena across the membrane, but also on most functions of the cell. In a recent modeling study^13^, we showed how deregulation of the dominant glial Kir current (*K*^+^ inward rectifier) could introduce strong nonlinearity into the total glial current, making the astrocyte prone to bistability switching when perturbed by a seizure-like local field potential (LFP).

In physiological conditions, astrocyte networks are electrically isopotential^5^, maintaining approximately the same *V*_*r*_ and being capable of rapidly dissipating voltage disturbances and restoring this state^14,15^. With ample evidence that neuro-glial networks undergo transient depolarizations and a wide range of local-field perturbations, it remains a challenge to describe the glial isopotentiality as a dynamical feature.

When astrocytic connectivity is assessed across the hierarchy of timescales it exploits, these networks appear to be rather plastic structures, both during development^16, 17^ and after reaching maturity^15^.

Although this study explores short time scales, i.e., the fast transient responses of *V* from tens of milliseconds to seconds, the slower molecular and structural changes offer rich parametric ranges to explore glial-specific, activity-dependent plasticity. (1) On a timescale of tens of seconds to minutes, GJ channel proteins undergo different phosphorylation events, depending on the signaling pathway. These changes dynamically alter open-channel probability, as shown for glial-specific GJ channels^18^. (2) On a slower scale, sustained phosphorylation of astrocytic GJCs has been shown to result in the dissociation of individual channels from the junction^19^.

As with most of the other astrocytic features, their connectivity also displays regional heterogeneity^20^, in line with cellular density and the structure–function demands of the local circuitry. As a result, syncytial coupling varies substantially from brain region to region, which can be precisely quantified by measuring syncytial isopotentiality. For example, compared with the hippocampus, syncytial isopotentiality is about 3.5 to 7.6-fold stronger in the visual, sensory, and motor cortical regions^14^.

In a wider context, the computational studies of neuro-glial network interaction today are the subject of growing focus, exploring a very wide range of structural and functional details; see Manninen et al. (2023)^21^ for a review. To what extent the synchrony within glial networks parallels the synchrony of neuronal networks is one of the central questions that quantitative neuroscience modeling is trying to address. A recent study has shown that glial regulation of extracellular *K*^+^ contributes to hippocampal synchrony specifically via the neuronal Kv7 channels, controlling afterhyperpolarization in neurons^22^.

In addition to the increasing focus on glial cells in all domains of neuroscience, dynamical systems models have not yet adequately represented glial networks. With an isolated astrocyte being only a building block of a real network, a proper quantitative description of glial coupling at the cellular level is needed to properly address electrical responses of glial clusters on timescales of milliseconds to seconds. To address this gap, in this study, we develop a minimal low-dimensional model of a coupled glial cell, incorporating both the voltage dependence of GJ conductance and the GJ activation kinetics of all junctions. Under common simplifying assumptions, we performed a limited parametric analysis on a “truncated” system— a self-coupled glial cell, representing an inner cell within a 1-D array of coupled glia, as the simplest cell-array model. Several bifurcation scenarios, presented in more detail, illustrate various forms of *V* multistability caused by coupling. Numerical simulations of a small 1-D array illustrate that nonlinear coupling impacts *V* dynamics, governing the propagation or failure of instabilities along the array.

## 2. Mathematical model of coupled glial cells

### 2.1 Electrical properties of isolated cell

In physiological conditions, the resting membrane voltage *V*_*r*_ of an uncoupled cell lies between −90 *mV* and −70 *mV*, near the *K*^+^ and *Cl*^−^ reverse potentials^11^, near which their respective channels are permeable. This range is close to, but 10 to 15 *mV* more negative than, the neuronal resting membrane voltage. That is the operating range of the key ion transporters, which engage the astrocytes in “housekeeping” of transient ionic imbalances following sustained neuronal activity. The stability of *V*_*r*_ is predominantly controlled by the ATP-driven *Na*^+^/*K*^+^ ion pump^23,24^ (NKA) through the maintenance of the Nernstian reverse potentials. The glial pump has different electrogenic properties from the neuronal variant^24,25^, which are significant for the differential role of astrocytes in ion homeostasis. In performing that role, in addition to the NKA pump, several leak-type *K*^+^ channels equip astrocytes with high *K*^+^permeability across a wide *V* range. Because the membrane is leaky near *V*_*r*_, mature astrocytes display very low input resistance, *R*_*m*_, typically between 2 and 5 *M*Ω in slices, as measured with single- or dual-electrode recordings^16^.

When present, nonlinearities in total astrocytic current *I*_*glia*_ arise from the nonlinear steady-state I–V profile of the major *K*^+^currents^13^ through Kir, K2P, and to a lesser extent other leak channels. Although Kir and K2P belong to the class of open, non-gated channels, their currents show complex electrochemical dependence. The Kirchhoff ODE equation for a node/current loop in a single compartment model takes the typical form:

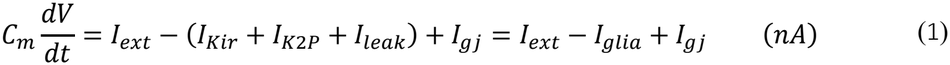

where *I*_*ext*_ is the external constant, or slowly varying current originating from permanently varying LFPs, while *I*_*gi*_ denotes the external current through all GJs of the cell. A striking electrophysiological feature of the astrocytic membrane is the nearly linear steady-state I–V curve of *I*_*glia*_, despite each of its individual additive components being nonlinear. Figure 2A (solid blue line) shows the nominal profile of the Kir current^26^, which is weakly inward-rectifying for positive *V*. Figure 2B shows how a sum of visibly rectifying currents results in an almost non-rectifying total current. In Fig. 2C, using the model of Janjic et al. (2023)^13^, we illustrate how a realistic, nonphysiologically “deformed” *I*_*Kir*_ profile produces an N-shaped *I*_*glia*_ (Fig. 2B), implicating instability of *V*_*r*_.

**Figure 2.**
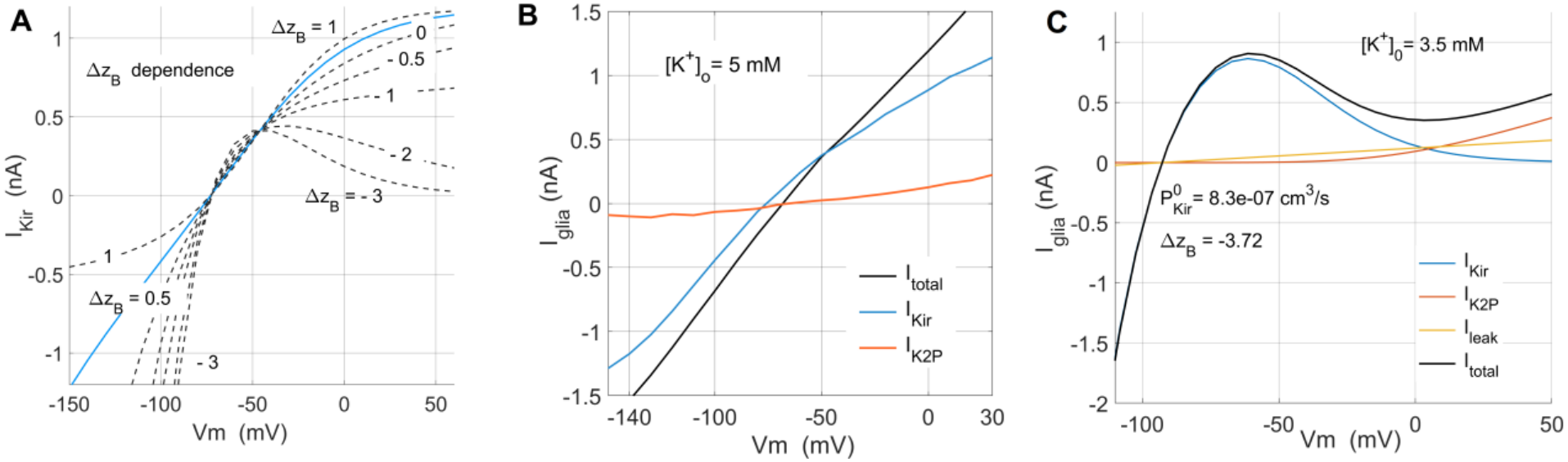
Model of nonlinear astrocytic currents. **(A)** Nonlinear Nernst–Planck description of *I*_*Kir*_, Eq. (2), shaped by Δ*z*_*B*_, the charge equivalent of modulatory channel block. The solid blue line, Δ*z*_*B*_ = 0.5 represents the nominal, physiological profile^13^. **(B)** Inward and outward rectified *I*_*Kir*_ and *I*_*K*2*P*_, respectively, from hippocampal glia, add up to non-rectified *I*_*total*_ .(A) and (B) from Janjic et al. (2023)^13^ with permission. (C) Example of N-shape deformed *I*_*glia*_ showing experimental deregulation turning weakly rectified *I*_*Kir*_ into strongly inwardly rectified, *z*_*B*_ < −1 with *I*_*K*2*P*_ and *I*_*leak*_ in their nominal profiles. *P*_*Kir*_in **(C)** is twice as large as in (A) to reflect astrocytic cluster effect on passive properties, rather than that of an isolated cell in (A). See *Supplementary Material S3*.

We model astrocytic currents using the Nernst–Planck description of an open pore, extended with thermodynamic formalism for pore activation.^13,27,28^ This description is extended with a Boltzmann activation term, describing the Coulombic, voltage-dependent, physiological block by positive polyvalent ions. The explicit form^13^ of *I*_*Kir*_ is given by:

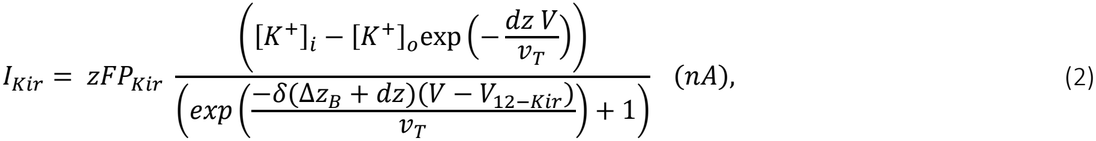

where *Δz*_*B*_ is the charge equivalent of the physiological block, and is the main parameter shaping *I*_*Kir*_ through *δ*(Δ*z*_*B*_ + *dz*). The later represents the resulting carrier charge^29^ that modulates the permeability of Kir channels. It controls the shape of *I*_*Kir*_ (*V*), Fig. 2A, and, in effect, the nature of rectification. Permeability *P*_*Kir*_ and *dz* produce additional shaping^13^; see *Supplementary Material S3*. The effective ion valence *dz* of the permeant *K*^+^ion(s) is one of the few fitted parameters, and could be *dz* > 1. The thermal voltage slope factor *v*_*T*_ ≡ *RT*⁄*F* equals 25.7mV at a room temperature of 25°*C*, with the universal gas constant *R* = 8.314 *J*/*mol K* and the Faraday’s constant *F* = 96485 *C*/*mol*.

The *I*_*kir*_ rectification, which produces an N-shaped *I*_*glia*_, introduces membrane bistability and switching^13^, as shown experimentally in cardiomyocytes with a nearly equivalent mixture of conductances^30^. The bistability, which manifests robustly over a wide range of *I*_*ext*_ and [*K*^+^]_*o*_, switches *V* between the moderately depolarized *V*_*r*_ and the very depolarized *V*_*dr*_ > −20*mV*, both of which are stable. The *V*_*dr*_ range represents catastrophic depolarization if it persists in real cells—a new equilibrium that would affect all voltage-controlled astrocytic functions. This possibility—that *V* alternates between *V*_*r*_ and the notably depolarized steady state *V*_*dr*_ illustrated in isolated cell model^13^, was the main motivation for extending that study to include nonlinear coupling. Although experimentally isolated *I*_*Kir*_ and *I*_*glia*_ have indicated nonlinearities that produce bistability, known experimental limitations associated with the leaky membrane would very likely have prevented detection of switching, even if the choice of parameters suggests that the phenomenon is reproducible^4^.

Even though they are much weaker, *I*_*K*2*P*_ and *I*_*leak*_ in Eq. (1) shape *I*_*glia*_ to the right of the *I*_*kir*_ inflection point, producing the N-shaped curve. In terms of kinetics, the smaller *I*_*K*2*P*_ shows *k*-order activation, *n*^*k*^, and turns on near −40*mV*, with *τ*_*K*2*P*_ ≈ 3 *ms*, which is still much slower than the momentary activation of *I*_*Kir*_ within 1 millisecond after the capacitive transient. As in the formulation of *I*_*kir*_, the extended Nernst-Planck formalism suggests *I*_*K*2*P*_ = *n*^*k*^*I*_*K*2*P*−*GHK*_, where *I*_*K*2*P*−*GHK*_ being the steady-state Goldman– Hodgkin–Katz current equation^31^ and *k* = 2. For the full expressions of the currents, and the values of the fixed parameters, see *Supplementary Material S5*. The minimal ODE model of an isolated astrocyte with constant ion concentrations is therefore given by^13^:

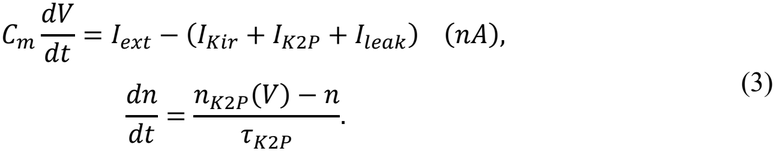

#### Gap junction conductance

Electrically, one GJ represents a highly conductive macroscopic connection^32^ that varies in the number of GJCs and in physical size, thereby creating an electrically distinct membrane domain (Fig. S1C, *Supplementary Material S1*). Most of the biophysical properties of GJ connections have been formulated based on two-electrode or dual patch recordings from two connected cells. Although this remains a standard protocol, the setup is limited because it cannot estimate the electrical and kinetic properties of a single GJ; see *Supplementary Material S2* for the equivalent circuit. This circuit records only from two terminals, summing up the contributions of all junctions into a single current readout that approximates the total conductance as if it arose from a single, lumped-parameter junction.

Single GJC conductance estimates range from 50 to 140 *pS* ^33–35^ for channels built by connexins of types 43 (Cx43) and 30 (Cx30), subtypes specific to glia^36^. Whole-cell studies based on native GJCs in astrocytes^5^ suggest a maximal conductance of 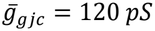 for a fully open GJC. Two gating states correspond to distinct permeation states^34,37^—a fully conductive state and a residual (partly conductive) state^38,39^, the latter estimated to be 20%–40% of the maximal conductance (Fig. 3A–3D). Based on this 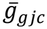 and an estimated average of 2,000 GJCs (of all junctions) between two neighboring astrocytes, estimated an upper bound of 240 nS for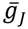, see Supplementary Information in Kiyoshi et al. (2018)^14^.

**Figure 3.**
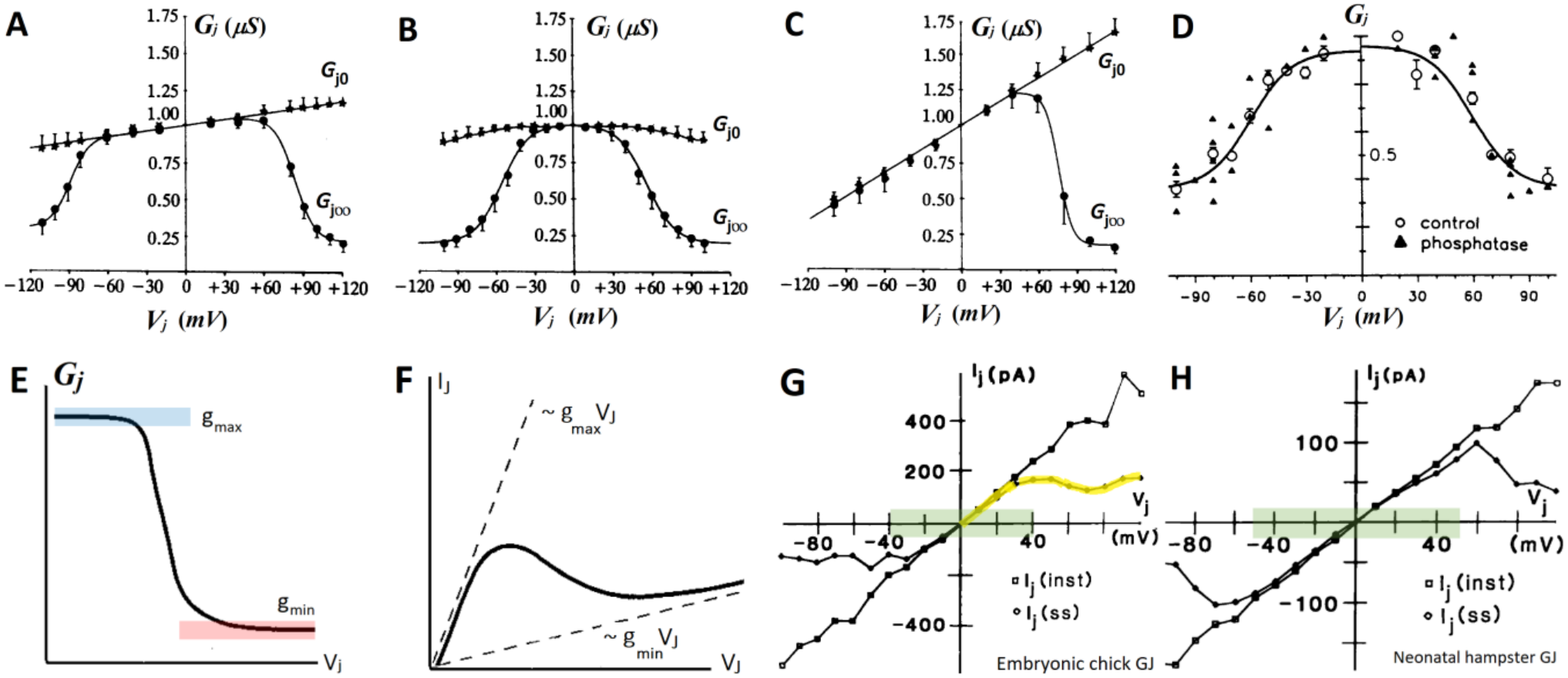
Instantaneous vs. steady-state voltage dependence of GJ conductance. **(A–C)** Voltage dependence of homotypic Cx26/Cx26, Cx32/Cx32, and heterotypic Cx26/Cx32 gap junctions, respectively, showing a marked difference between the instantaneous *G*_*j*0_ (filled asterisks) and steady-state conductance *G*_*j*∞_, (filled circles). Figures modified from Barrio (1991),^42^ with permission. **(D)** Normalized *G*_*J*_ − *V*_*J*_ dependence of GJs formed by Cx43 gap junctions, showing a high normalized residual conductance, *g*_*min*_, characteristic of astrocytic Cx43 junctions. Reproduced from Moreno (1992)^43^, with permission. All steady-state dependences in (A–D) are fitted with the Boltzmann equation for first-order inactivation kinetics, with residual conductance *g*_*min*_. **(E)** A sketch of a single-sided Boltzmann relation with a residual term, approximating the profiles in (B) and (D) and producing the N-shaped *I*_*J*_- *V*_*J*_ current profile in (F). Sketches modified from Baigent (2001)^44^, with permission. **(G, H)** Instantaneous (squares) and steady-state (rhombs) currents through gap junctions from chick embryos and neonatal hamsters, closely reflecting the qualitative shape of the voltage dependence of conductances in (A–D). Modified from Veenstra (1992)^35^; permission requested from T&F via form. Within the *V*_*J*_ range of *G*_*J*_ plateau in (A–D) (shaded green), the measured *I*_*J*_ shows the expected ohmic profile. The positive branch of the *I*_*J*_ current in (G), marked in yellow, matches the N-shaped profile sketched in (F).

#### Voltage dependence and its kinetics of the gap junction conductance

In what follows, we refer to potential difference *V*_*J*_ = *V*_*B*_ − *V*_*A*_ as transjunctional voltage between the interiors of cells A and B. Voltage-dependent GJ inactivation with increasing absolute *V*_*J*_ is the main GJ kinetic change driven by voltage. Most junctions formed by homotypic GJCs (built by a single Connexin homolog) remain open over a wide range, near 60 mV, of instantaneous *V*_*J*_. They partly inactivate, reducing their conductance to a lower residual level under prolonged polarization^40^. In the rest of the chapter, we describe GJ properties using their dependence on *V*_*J*_ rather than *V*, as is typical in experimental studies (Fig. 3A–3H). The *V*_*J*_ differences encountered later in the stability and bifurcation analysis, *V*_*J*_ = *V*_*dr*_ − *V* or *V*_*J*_ = *V*_*r*_ − *V*, with constant *V*_*r*_ and *V*_*dr*_, make *V*_*J*_ only a translational shift in the coordinate, without other algebraic effects. Figure 4A–4C shows experimental data of *g*_*J*_ inactivation for GJCs formed from different connexin combinations. Recorded junctional currents, *I*_*J*_ (Fig. 3), sum up the currents through all GJ connections between two cells^32,41^, suggesting a single value of *τ*_*J*_. We therefore model GJ connection properties as arising from a sum of junctions with an average number of channels and an average plaque size^32^.

**Figure 4.**
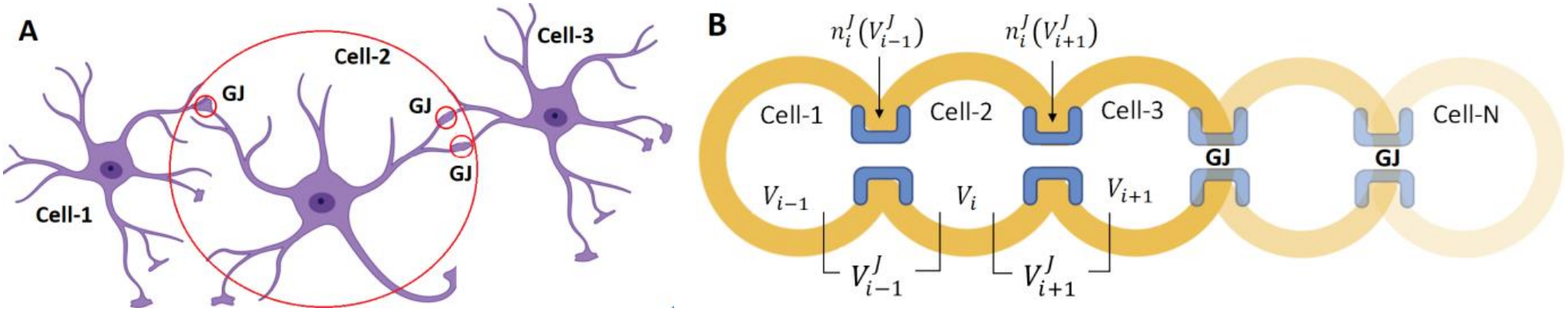
Arrays of connected astrocytes. **(A)** Astrocytes are connected by a variable number of GJ connections, each formed by a cluster containing a variable number of channels, mostly, though not exclusively, at the tips of branched processes. Each cell spatially occupies one astrocytic domain^50^, outlined in red. **(B)** One-dimensional array model of glia, in which the total number of GJs and their channels is divided between the left and right connections. Although biologically simplified, the model still captures the key aspect of the different responses of junctional voltages 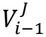 and 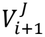 to asymmetric perturbations originating from one side. It also describes a common feature: a subset of GJs at the left connection is perturbed before those at the right, generating very different initial voltages 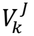. Both drawings were created in *BioRender*.

#### Gap junction inactivation kinetics

To detail the kinetics of the macroscopic junctional current, we use the Hodgkin–Huxley term 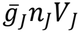. For the GJ inactivation probability *n*_*J*_, we suggest first-order kinetics with a time constant *τ*_*J*_, similar to those proposed in early studies of coupled amphibian cells^45,46^. Even though this approach notably simplifies the gating complexity of GJCs^37,47,48^, it is sufficient to capture the single-sided behavior of *g*_*J*_(*V*_*J*_) as shown in Fig. 3E, given the symmetry of the *V*_*J*_ profile. For *n*_*J*_ and the steady-state voltage dependence 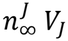, we get:

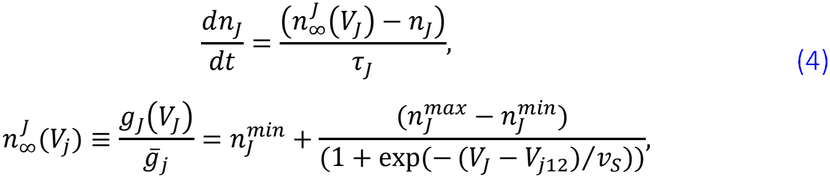

where 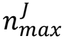 and 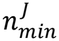 are open channel probabilities corresponding to the maximal and residual conductance of all GJs between two cells, *V*_*j*12_ is the half-inactivation voltage, while the positive parameter *v*_*S*_ is a voltage slope factor of the sigmoid, to be later tuned to Cx43 junctions (Fig. 3D). The non-zero residual probability 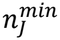 of Cx43 junctions is the key difference from the single, open–closed-gate Boltzmann equation. It has a strong impact on the *V*_*r*_ stability analysis, as illustrated later. The form of Eq. (4) ensures that the normative range of 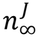 stays below unity, between 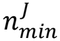 and 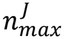 . With *V*_*j*12_ between 30 and 50 mV, 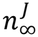 reduces to 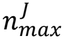 for fully conductive GJs near *V*_*J*_ ≈ 0.

Without data on kinetic variability between small and large GJs, the whole-cell estimates of *τ*_*J*_ in dual-patch recordings^32,45^, between 70 and 200 ms, represent averages over all junctions. Instantaneous currents reach their peak order of magnitude faster, in less than 10 ms^4,14,15^. The timescale separation between *τ*_*J*_ and *τ*_*K*2*P*_ in Eq. (3) as well as the separation between *τ*_*J*_ and the membrane time constant *τ*_*m*_, which are one to two orders of magnitude smaller than *τ*_*J*_, suggests treating K2P activation as instantaneous relative to junctional inactivation with time constant *τ*_*J*_, and omitting K2P activation from Eq. (3). A constant *τ*_*J*_ = 100 *ms* ms is used throughout the simulations.

### 2.2 1-dimensional array model of connected astrocytes

In addition to being structurally described^6^, the complex 3-dimensional coupling of mammalian astrocytes has not yet been quantified across a larger sample, leaving us without estimates of the average number of junctions per peripheral process or branch and per cell, or of the distribution of the number GJC across the junctions. The spatial organization of mammalian astrocytes, which occupy non-overlapping volumetric, 3-D domains^49,50^ in some brain areas and are interconnected by GJs at their tips (which define domain boundaries), makes such quantification critically important for formulating meaningful models.

Although attempts have been made to model and simulate interconnected glia, most models that explicitly model voltage lack biophysical detail about astrocyte connectivity because of their different research focus; see Manninen et al. (2023)^21^ for further references. For models focused on Ca^2+^ dynamics and intercellular calcium waves, see Goldberg et al. (2010)^64^ and Matrosov et al. (2019)^65^.

Notation. Within the array formalism, lowercase “*i*” in subscript denotes array indexing, while uppercase “*J*” in the superscript labels a junctional property. Within single-cell ODEs, the junctional index is instead written as a subscript.

The simplest non-biological constructs traditionally used to model coupled cells are 1-D arrays of *N* single-cell ODEs coupled by GJs. Although a 1-D array drastically simplifies complex 3-D connectivity to left and right connections, it suffices to illustrate the effects of coupling on *V*_*r*_ stability. We begin with the basic ODE of an inner cell:

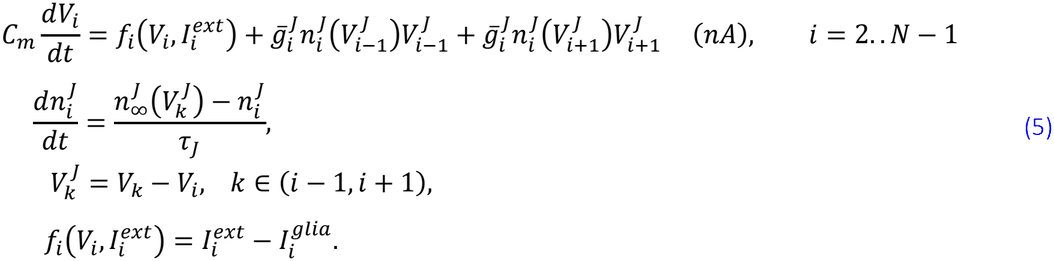

We emphasize that the junctional voltage 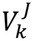 has different values at different junctions of *i*-th cell; in the array notation, it is defined as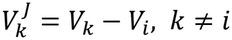. Here, we assume that the junctions are homotypic and have (i) symmetric conductance, (ii) the same kinetics, and (iii) connection strengths that sum to half the total across all junctions (Fig. 4B).

Equation (5) represents the *i*-th inner cell in an array model of discretely coupled, second-order reaction– diffusion ODEs, as used in many multicellular systems^51,52^. However, it does not yet constitute a proper ODE model, since it does not account for the first and last cells, it requires 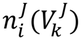 to be evaluated twice^1^ (one for each junction), and the boundary conditions have not yet been specified. We will address these issues by further assumptions in the subsequent development of the model.

#### Single active site

The multistability of *V*_*r*_ will be explored in an inner, *i*-th cell described by Eq. (5), with the (*i*-1)th cell depolarized and kept at *V*_*dr*_, while the next cell, the (*i*+1)th cell, and the remaining cells are initially kept at *V*_*r*_. With the first cell perturbed, the steady-state stability and bifurcations of the second cell (Cell 2) are central to the study. This approach of using a *single active site* has already been used in solitary wave and front propagation models in discrete bistable media^*53,54*^. For simplicity, we will keep the same resting state *V*_*r*_ in all cells, as indicated earlier. The propagation effects depend on (a) the separation of the fixed points, usually analyzed using the saddle quantity *a* = |*V*_*r*_ − *V*_*s*_|/|*V*_*dr*_ − *V*_*r*_|^54,55^, and (b) the slow inactivation governed by *n*_*J*_. The large timescale separation warrants this reductionist approach, which makes bifurcation analysis manageable. With simplified *f*, asymptotic analysis using perturbation theory may provide criteria for front propagation^55^.

### 2.3 Reduction to a minimal model

The much slower inactivation 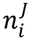 compared with the membrane time constant (implicit in *f*), *τ*_*m*_ ≪ *τ*_*J*_, allow 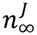 to be approximated by a constant term for small *V*_*J*_ (Fig. 3E and Eq. (4)). This, in turn, allows Eq. (5) to be rewritten for an inner active site, with only a single evaluation of *n*_*J*_, thereby making Eq. (6) a proper ODE that can be used in numerical phase-plane and bifurcation analysis:

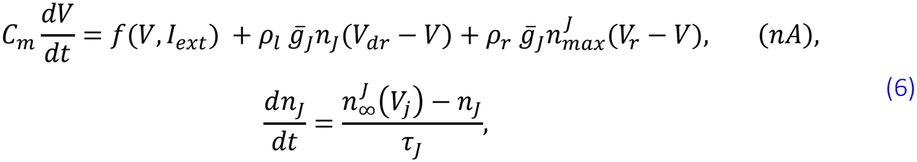

where, for clarity, we have omitted the array indexing and written an explicit expression for each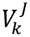. The following assumptions allow us to obtain Eq. (6). **(1)** With (*i*-1)-th cell already depolarized, the left connection of the (*i*)-th cell experiences a large positive voltage difference, 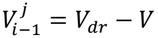, within the transition region of *g*_*J*_(*V*_*J*_) (Fig. 3E and Eq. (4)); hence, it requires evaluating 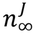 which governs the nonlinearity in *I*_*J*_(*V*_*J*_) (Fig. 3F). **(2)** By contrast, the (*i*+1)th junction to a quiet cell at *V*_*r*_, experiences a small voltage difference, 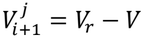, near zero. Its GJs remain fully open, with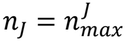, over a broad range of *V*_*j*_, for transients lasting at least *τ*_*J*_. This allows us to assign a linear, ohmic current profile *I*_*J*_(*V*_*j*_) = *g*_*max*_*V*_*J*_ to that junction. This simplification requires that, with Cell 1 at *V*_*dr*_, the active site, Cell 2 is not already strongly depolarized and remains very close to the saddle *V*_*s*_. This is equivalent to requiring the saddle quantity to satisfy *a* ≫ 0, or that *V*_*r*_ be notably separated from the saddle.

Assigning unequal fractional multipliers *ρ*_*l*_ and *ρ*_*r*_ to the junctional current terms in Eq. (6) reflects the fact that each astrocyte connects to approximately ten neighbors and exhibits a diffuse rather than directional spread of depolarization. Any cell near a propagating boundary is engaged by multiple neighboring cells. This volume effect means that, at any given moment, an inner cell has more GJs engaged than remain associated with quiet cells, implying that scaling with *ρ*_*l*_ > *ρ*_*r*_ brings the simplified coupling model closer to reality. In the numerical bifurcation analyses, we kept the scaling fractions close to *ρ*_*l*_ = 0.8, and *ρ*_*r*_ = 0.2.

#### Self-coupled cell

Equation (6) is our minimal model suitable for numerical bifurcation analysis. Within the class of simple models, it represents a variant of the Bonhoeffer–Van der Pol system. It describes a single self-coupled cell with *f* fulfilling the non-monotonicity condition for bistability of *f* ^56^. The additional condition of bounded perturbations^56^ is *a priori* granted in biological settings by finite and observable perturbations *giV*_*j*_ and *I*_*ext*_. Various forms of such nonlinear negative-feedback models with sigmoid activation arise when modeling neuronal inhibitory autaptic connections (auto-synapsing).

The ranges of some passive parameters, such as maximal conductances, input resistance, or total membrane capacitance, need numerical adjustments when estimated from recordings of an isolated cell, in which they are, in most cases, underestimated. Recordings from intact tissue always pick up a signal from an “effective cell”—an irregular assembly formed by the soma of the patched (measured) astrocyte and parts (peripheral astrocytic processes) of several strongly connected immediate neighbors, due to the very conductive GJs under nominal physiological conditions^4,5,14^ (see *Supplementary Material S3)*.

### 2.4 Numerical methods

Basic stability analysis and exploratory numerical integrations of ODEs in Eq. (1) and Eq. (5) were performed in *XPPAUT* ^57^, while all bifurcation analysis was performed using *AUTO*^*58*^ integrated within *XPPAUT*. The default ODE solver used in *XPPAUT* is the fourth-order Runge–Kutta scheme, with a fixed time step in the range of 0.02 – 0.05 ms. Time steps of 0.01 or 0.02 ms were used for the ODEs during continuation in *AUTO*. The typical initial continuation step in AUTO was ±0.01. Sensitivity near some peculiar points, such as the limit points of cycles (see below), required starting with 0.005. Such boundary regions were also validated using *MATCONT*^*59*^, as implemented within the MATLAB-based *MATCONT Toolbox*. We comment in Chapter 3, “Numerical bifurcation analysis and simulations,” on a case where the two continuations differed.

Numerical integration of the array model, Eq. (5), was performed in *MATLAB*, using Newton’s method with a time step of 0.01 ms, as it was the most efficient method. *MATLAB*’s *ode23* solver, with an adaptive step size, was used for validation where higher sensitivity was noticed, but it never resulted in a different solution type or visibly different transients. *Ode23* is an implementation of a medium-accuracy, non-stiff ordinary differential equation solver that uses explicit second- and third-order Runge–Kutta formulas. The array was integrated in parallel, with a free boundary condition at the *N*-th cell and other initial conditions as described above.

## 3. Numerical bifurcation analysis and simulations

As indicated by the feature analysis of GJ coupling, in addition to the instabilities introduced by nonlinear *f*, increased bifurcation complexity should be expected due to the nonlinear coupling. The SN bifurcation of *V*_*r*_ with increasing *I*_*ext*_ in the self-coupled system of Eq. (6), still kept in ℝ^2^, has a central place. All neural cells respond to physiological depolarizations due to *I*_*ext*_ as a principal perturbation triggering their transient responses. In that regard, it is not surprising that the effects of coupling we have illustrated here represent additional complexity brought by GJ coupling.

NOTE: The values of parameters of *f*(*V, I*_*ext*_) varied within the numerical bifurcation analysis are given in the corresponding sections. Parameter ranges for all other parameters, as well as the equations describing the astrocytic currents, have been reused from the published with the isolated-cell model^13^ and are listed in *Supplementary Material S7*.

### 3.1 Scenario 1: Saddle-node bifurcation of *I*_*ext*_ with a fold of limit cycles window

A fold bifurcation of *V*_*r*_ over *I*_*ext*_ is presented for Cell 2, with the neighbor at resting states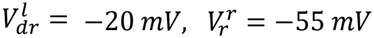, respectively. The fold structure was obtained with *V* continuation for 0.5 < *I*_*ext*_ < 0.6 *nA*.

The SN structure over *I*_*ext*_, generic in the isolated cell^13^, was altered by a narrow window of unstable periodic orbit (UPO) cycles emerging from the Hopf point HB1 on the stable *V*_*r*_-branch (Fig. 5A) for *I*_*ext*_ = 0.531126 *nA* and disappearing at the limit point of cycles (LPC; Fig. 5B).^2^ The critical biophysical effect introduced by the coupling is the major shrinkage of the stable, physiological branch of *V*_*r*_ by HB1, making *V* more prone to instability driven by *I*_*ext*_, compared with the isolated cell.

**Figure 5.**
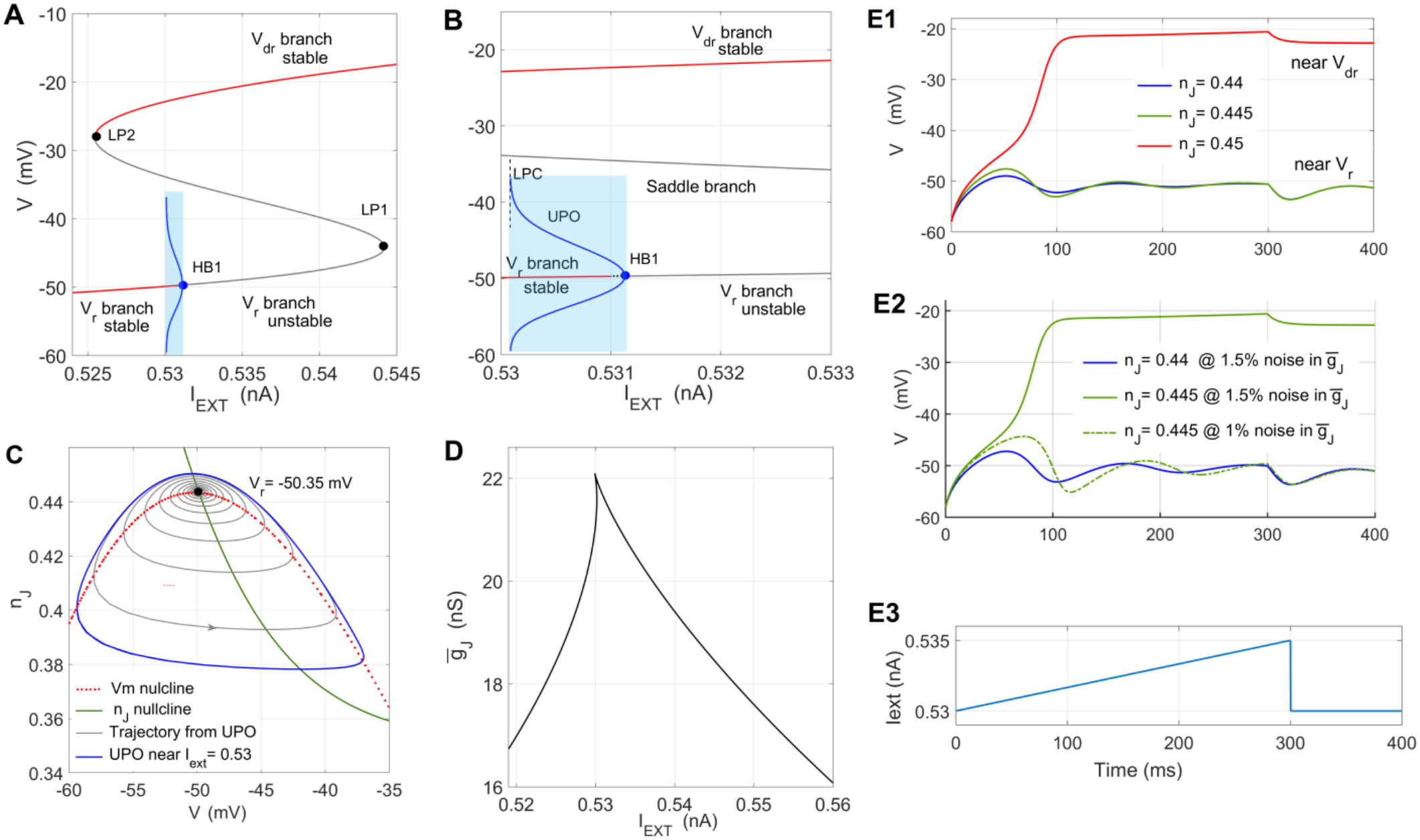
Saddle-node structure with an UPO window. **(A)** The generic fold-structure over *I*_*ext*_ with a Hopf point HB1 and an nstable periodic orbit (UPO) window, shaded blue. **(B)** The UPO window (0.53, 0.531) enlarged, with the symmetrical blue urve outlining the amplitude of the UPO. This structure shortens notably the lower stable branch of *V*_*r*_ (red branch). The ray segments in (A) and (B) represent the unstable branches of the fold curve. **(C)** A phase-space view of the UPO (blue) ear *I*_*ext*_ = 0.53, around a stable focus at *V*_*r*_ = −50.35 *mV*, with a period of approximately *T*_*UPO*_ = 303.8 *ms*. The gray rajectory starting at the UPO spirals inward to *V*_*r*_ . **(D)** A cusp curve over the same *I*_*ext*_ range indicates a rather wide range of 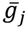, itself producing a fold around *V*_*r*_ . (E1–E3) Noise-driven bistable switching near the Hopf point. (E1) Examples of three rajectories starting near (−58 *mV*, 0.44) in the (*V, n*_*J*_) plane, with *V* depolarized by the ramp in (E3). With *V*_0_ = −58 *mV*,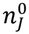 moves the initial state closer to or farther from the UPO (see panel 5C). For *n*^0^ values inside the UPO, the green and blue ransients take the initial state to *V*_*r*_ . For 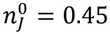, the red curve shows a switching transient *V*_*r*_ → *V*_*dr*_ . (E2) Same as in (E1), ith up to 1.5% noise added to 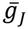. Already for 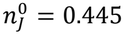 (green graph), the noisy conductance takes the state to *V*_*dr*_, hich was not the case in (E1). To show that this sensitivity comes from the noise, the green dash-dotted line shows the ame solution with the noise amplitude decreased to 1%, which does not trigger switching to *V*_*dr*_ . (E3) The ramp profile of he *I*_*ext*_ current used to depolarize the cell in (E1) and (E2), with *I*_*ext*_ in the blue-shaded region. Locating 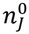 values and rajectories in (C) is not possible due to the ramp-induced change in *I*_*ext*_, which shifts the nullclines, moving the steady states nd UPO. Fixed parameter values (Fig. 5): 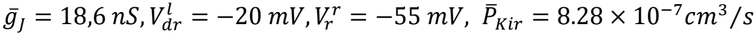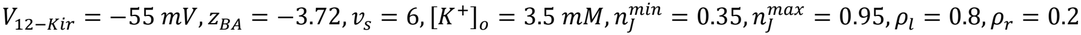.

A phase-plane example of the UPO is shown in Fig. 5C, with an approximate period of *T*_*upo*_ = 303.8 *ms*, for the parameters given in the Fig. 5 caption. To visualize the 3-D geometry of the thin repulsion UPO layer in ℝ^2^ × ℂ^1^, see Izhikevich (2007, Chapter 6.1.4)^60^. The UPO amplitudes saturate at the boundary of the narrow layer, representing the limit point of cycles (LPC point; Fig. 5B). A typical behavior of *V* near the UPO is a pseudo-periodic transient ending at *V*_*r*_ or *V*_*dr*_. Figure 5C shows an inward bound transient ending at *V*_*r*_.

The fold structure is generic when varying 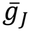 as well, shown by the cusp diagram in the *I*_*ext*_, 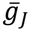 plane (Fig. 5D). This case lies near the 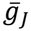 branch of the cusp.

Any variant of a slow change in *I*_*ext*_ through HB1, mimicking an incoming depolarization wave, will provide a sufficiently long dwell time for the state near HB1 to allow *V* to be repelled by the UPO. The examples in Fig. 5E1–5E3 illustrate this effect: parameter noise added to 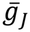 produces switching near the UPO. A ramp profile of *I*_*ext*_ takes the state in the blue-shaded region of Fig. 5A and 5B, where, for fixed *V*_0_ = 58 *mV*, slightly varying 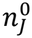 initiates transients starting near the UPO. Parameter noise of up to 0.3 nS (1.5%) was added to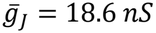, using normally distributed random numbers. It is neurobiologically important to highlight that, in Fig. 5E2 and 5E3, after the *I*_*ext*_ ramp ends, *V* still dwells near the non-physiological, but stable *V*_*dr*_. This shows pronounced system sensitivity associated with the shortened stable branch in Fig. 5A, due to the UPO band. Switching occurs somewhere after 50–60 ms, where 0.53 < *I*_*ext*_ < 0.531 (Fig. 5E3), placing the system within the UPO band and explaining the waviness of *V* near *V*_*r*_, which is absent near *V*_*dr*_.

The results of numerical simulations of a 1-D array, *N* = 4, are presented in Figure 6. For constant *I*_*ext*_ = 0.534 *nA*, the coupling strength was varied within 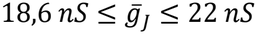. Color-coded temporal evolution of *V*_*m*_, total current *I*_*glia*_, and GJ inactivation *n*_*J*_ is shown for 10 seconds. The array is perturbed by keeping Cell 1 permanently at the depolarized steady-state *V*_*dr*_ = −20 *mV*, with a bistability configuration in *f*.

**Figure 6.**
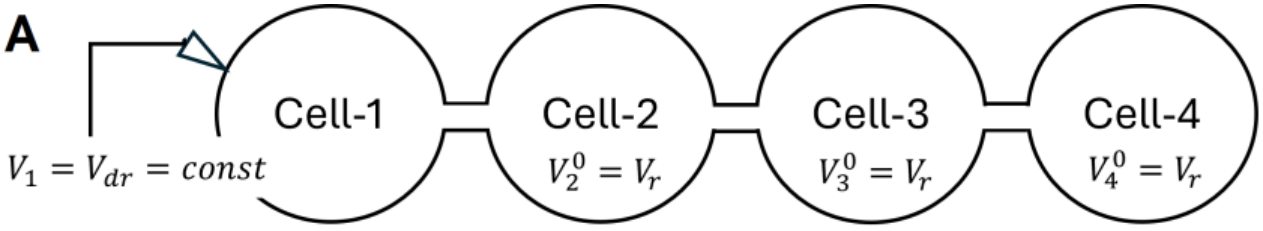
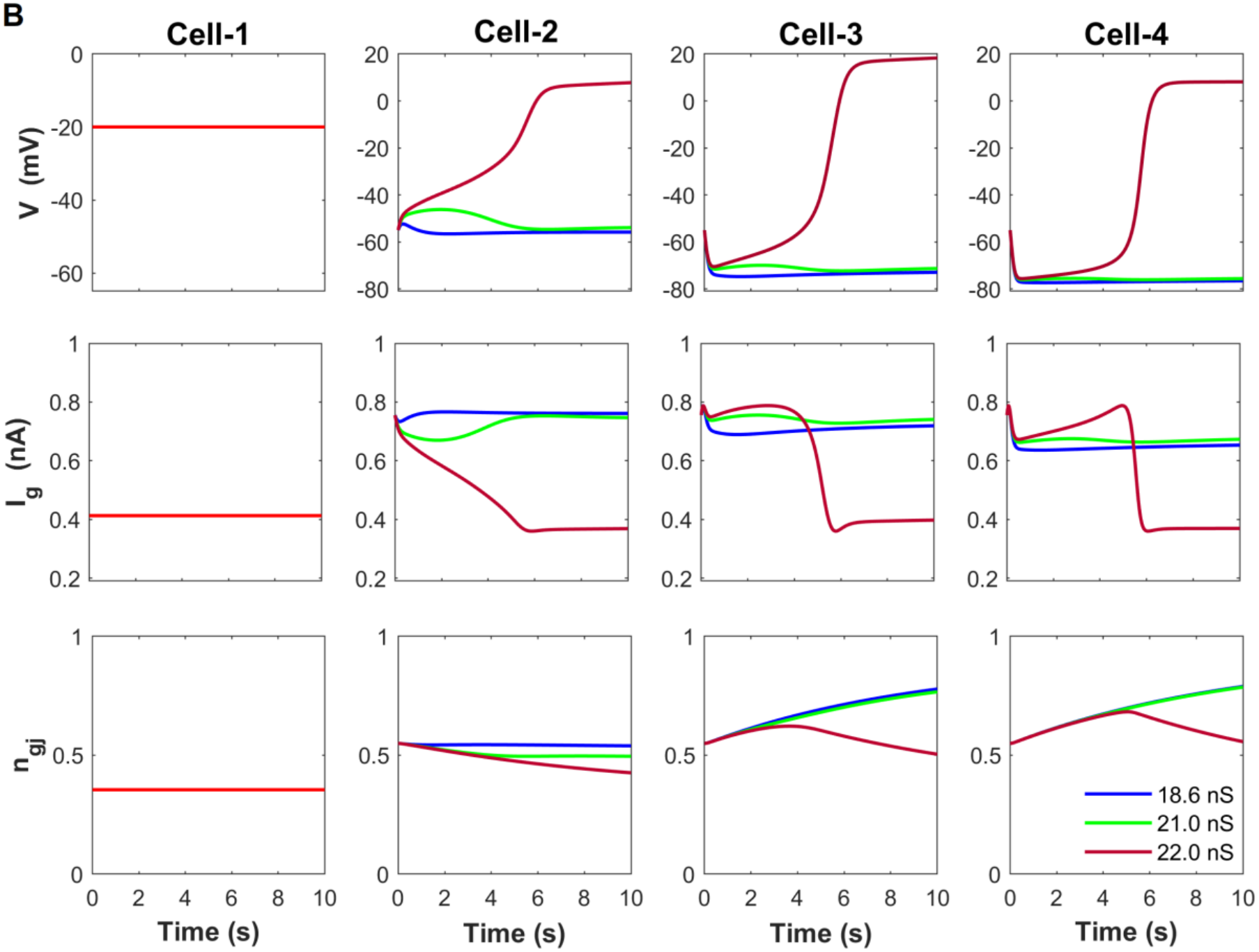
Temporal evolution of a 1-D array of N=4 coupled cells in Scenario 1. **(A)** Schematics of the numerical simulation of a 4-cell array, with Cell 1 under constant depolarization at *V*_*dr*_ = −20 *mV*, serving as a fixed boundary condition and a depolarized front perturbation. (B) Array simulation for *I*_*ext*_ = 0.534 *nA* and three values of the slope GJ conductance, 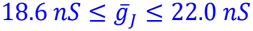, showing the transition to front propagation towards *V*_*dr*_ along the array. The color legend in the lower right panel applies to all transitions in Cell 2–Cell 4.

The transient behavior of array-connected cells shows two biophysically important details. First, we can validate the key assumption for qualifying the self-coupled cell as the target system, given in Eq. (7a) and (7b): the large time-scale separation due to *τ*_*J*_ ≫ *τ*_*m*_. The *n*_*J*_ changes are slow within the rather narrow dynamical range between 0.55 and 0.75, keeping the junctions conductive. Second, we see that stronger coupling produces greater *V*_*m*_ depolarization and stronger *I*_*glia*_ transients (red traces) for a still moderate *I*_*ext*_. This is not intuitive for Cell 3 and further down the array, since *V*_*J*_ polarity changes and the junctions are bidirectionally conductive.

### 3.2 Scenario 2: Enriched multistability with limit cycle bifurcations

The previous Scenario 1 illustrated that SN of *f* in an isolated cell, over *I*_*ext*_, is also present in the self-coupled cell. The following bifurcation scenarios show that maximal GJ conductance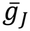, as the sole bifurcation parameter, enriches the SN structure, giving birth to stable periodic spiking over a measurable range of 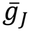 variation. The underlying fold structure was obtained with *V*_*r*_ continuation for 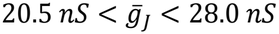. The new structure in this scenario appeared after lowering the residual open probability of GJs to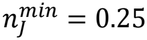.

Figure 7A describes the emergence of stable oscillations over a measurable, 5 *nS* wide interval of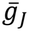. They appear by decreasing 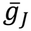, via a supercritical Hopf bifurcation at *HB*_*right*_, near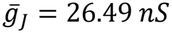, with the parabola (green arcs) extending toward lower values, ending in a narrow crisis window (Fig. 7A, blue shading).

**Figure 7.**
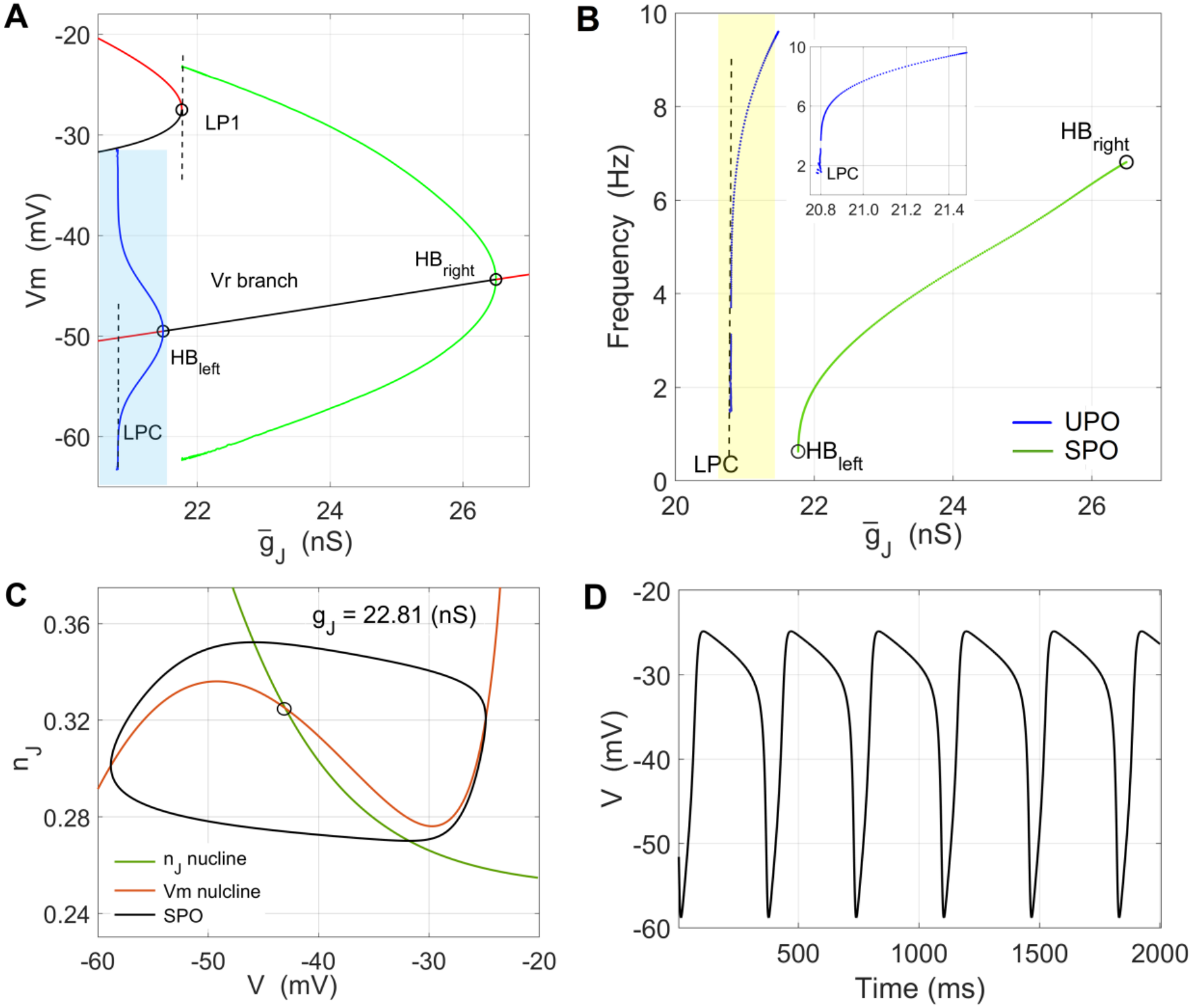
Large-orbit SPO meeting a “crisis” band of bistable behavior. **(A)** Bifurcation diagram in 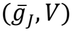 showing a 5 *nS* wide window of 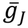, where SPOs emerge through a supercritical Hopf bifurcation as 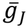 decreases along the *V*_*r*_ branch, with the green curve extending to the left of *HB*_*right*_. At LP1, this large-amplitude SPO disappears at the bistable switching window of the fold structure of the *V*_*dr*_ branch. The red segment represents a stable branch, while the gray segment represents the unstable branch of a fold curve. Blue segments designate Hopf curves of UPO onset, with blue shading marking their parameter windows. **(B)** Frequency plot of both intervals of limit cycles: of the SPOs in green and the UPOs in blue. **(C)** and **(D)** Phase-plane and time-series examples of a large-orbit SPO for 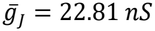 . Other changeable parameter values for (A–C) are as follows: 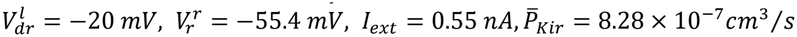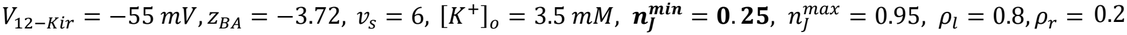.

A structure similar to that in Scenario1, Figs. 5A and 5B, is embedded within the fold structure as a band of UPOs less than 1 *nS* wide along 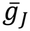, interrupting the *V*_*r*_ branch and creating an unstable segment to the left of *HB*_*left*_, at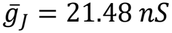. To the left, the UPO band extends up to the LPC at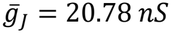.

The frequency plot in Fig. 7B shows the frequency dependence of both periodic structures. The green curve displays the known dependence for a supercritical Hopf bifurcation, with the frequency decreasing but staying finite as 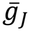 approaches *HB*_*left*_. The continuation of the Hopf curve ends slightly before *HB*_*left*_, at the LP1 boundary of the fold curve. With both points, *HB*_*left*_ and LP1, representing the boundary between stable and unstable behavior, the numerical sensitivity prevents further continuation. The blue curve in Fig. 7B describes the fold of UPO limit cycles, with the yellow-shaded area enlarged in the inset, showing frequency scaling according to 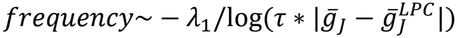. The frequency decreases but remains nonzero as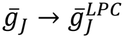, which is known to be a feature of a saddle homoclinic bifurcation (SHO)^60^, which we did not detect numerically. See Fig. S4 in *Supplementary Material 4* for the least-squares curve fitting.

Slight single-parameter variations in *I*_*Kir*_ or the coupling kinetics 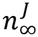, such as variations in *V*_*J*12_ or *v*_*S*_, result in different stable periodic structures within the same 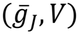 domain. See *Supplementary Material S5* for a case of a small-amplitude back-to-back SPO bifurcation structure within the (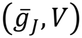) domain of the previous bifurcation diagram.

The enriched fold structure over a measurable range of 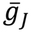 suggesting that weakening the coupling, or gradual uncoupling of the cells, may bring us into the range of periodic oscillations, is an exciting new observation in dynamical studies of the astrocytic membrane. Even though we analyzed a self-coupled cell here, it is a novel possibility in the fast *V* response of a model for which the assumptions underlying the reduced model Eq. (6) are valid.

We simulated the 4-cell array using parameters of the first of the two SPO scenarios, with *I*_*ext*_ = 0.55 *nA* and 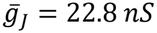 in Figure 8, corresponding to the large-orbit SPOs. The observed behavior for slightly increased 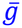 and an increased kinetic range of GJ inactivation (decreased 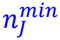) was a very fast jump from *V*_*r*_ → *V*_*dr*_, suggesting that the introduced coupling variation moves the state of Cell 02 into the bistability region, within the blue-shaded area in Fig. 7A. This could explain the more pronounced bistability with fast jumps, for the same fixed boundaries, 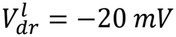 and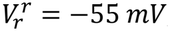, as in the first SN scenario (Scenario 1).

**Figure 8.**
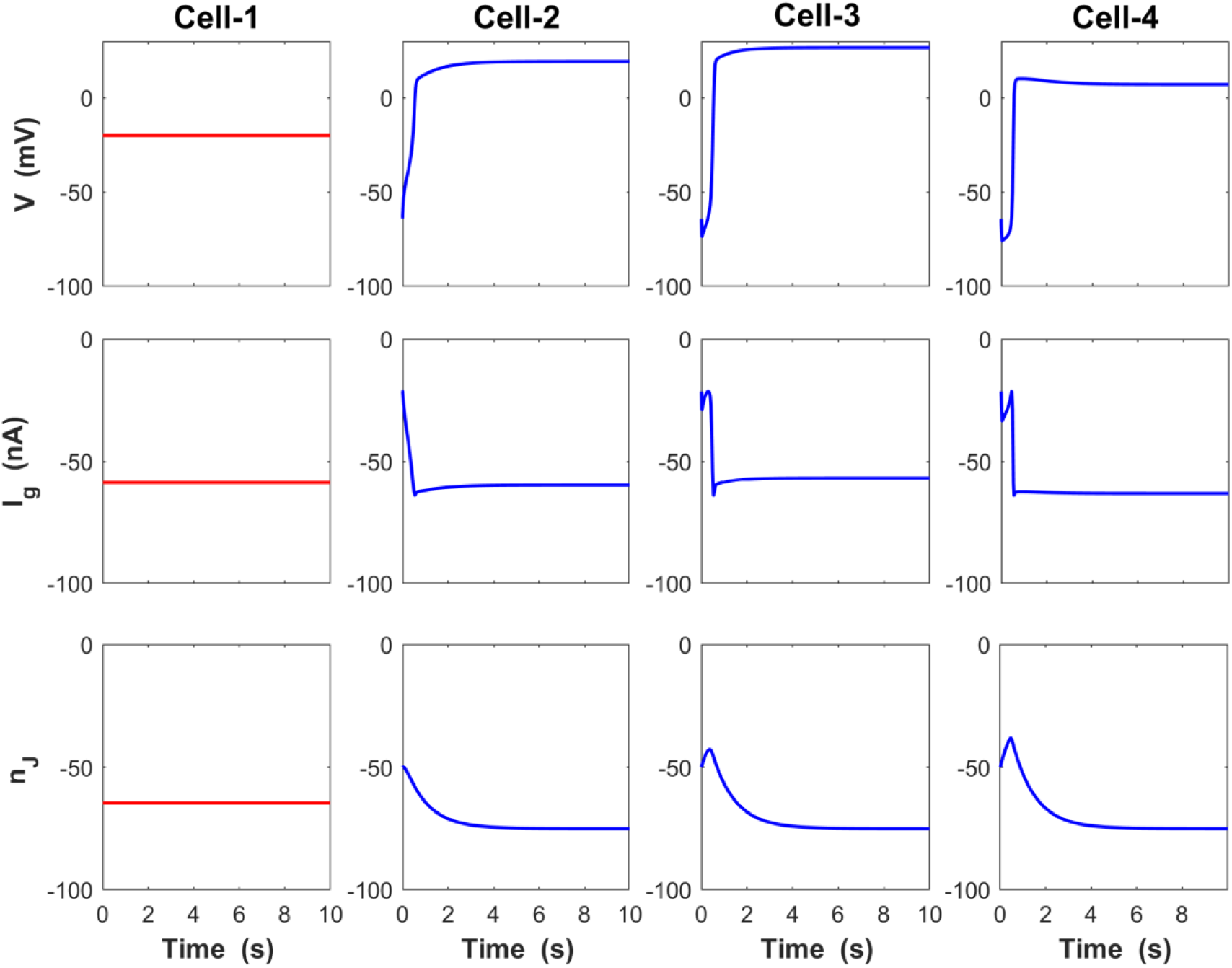
Temporal evolution of 1-D array of N=4 coupled cells in Scenario 2. Array simulation for 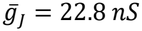 and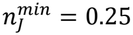, near *I*_*ext*_ = 0.55 *nA*, within the range of the SPO, close to Scenario 1. It shows a very fast jump transition to *V*_*dr*_ throughout the array. Cell 1 was kept on *V*_*dr*_ as a fixed boundary condition, serving as an array perturbation. The schematic of this array simulation is the same as the one in Scenario 1, Fig. 6A.

We stress here that 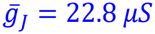 is close but does not fully match the corresponding parametric region in cusp bifurcation diagram (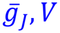) of self-coupled Cell 2 in Fig. 5D. In addition to altered *I*_*ext*_ and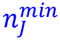, resulting in altered bifurcation branches, the coupling to a “live” Cell 3 makes the Cell 2 ODE non-stationary, with a shifting boundary 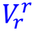,at the right GJ connection.

Array simulations suggest that constraining the array solution within the region of even transiently oscillatory solutions requires different strategies. Exploring multiple-parameter ranges to eventually arrive at criteria for observing periodic solutions in 1-D array is a complex challenge, beyond the scope of this study. The principal obstacle is the lack of coherent recordings of astrocytic currents and GJ properties in the same set of cells with nonlinear *I*_*Kir*_, such that at least parameter trends can be detected. Numerical bifurcation analysis in such an extended study will likely involve two-parameter bifurcations, lying close to those presented in the Scenario2 scenarios.

## Discussion

Experimentally, in physiological conditions astrocytic networks undergo small variations in *V*, which are attenuated by the network to restore syncytial or network isopotentiality^14,15^. Such behavior relies on highly conductive junctions for monovalent ions and a near-linear *V* dependence of the total astrocytic current. With growing evidence that deregulated glial Kir4.1 current is linked to neurodegenerative^61–63,66^, we found it important to extend the model of an isolated cell^13^ to coupled astrocytes, even though the bistability switching remains to be experimentally validated.

In this study, we went beyond GJ flux models using the leaky ohmic or Goldman–Hodgkin–Katz equation for 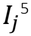 by introducing voltage and time dependence through the inactivation probability *n*_*J*_, an approach analyzed for the first time in glia using a self-coupled single-cell model. Although non-biological, we argued that a single cell coupled to fixed voltages is a relevant model for studying *V*_*r*_ instabilities over short timescales. Even though introduced decades ago, nonlinear *n*_*J*_(*V*_*J*_) kinetics of glial-specific Cx43 gap junctions, as described in Eq. (4), have not been introduced into astrocyte network models. We stress here that nonlinear forms of flux coupling mediated by glial second-messenger pathways, such as inositol trisphosphate (IP3) in the mediation of intra- and intercellular calcium waves^64,65^, are outside the scope of the present study, as they are not directly modulated by *V*_*j*_.

A key insight of this study is the demonstration that coupling further promotes bistability in the coupled cell model compared with the isolated cell model, with more bifurcation scenarios. The cusp in 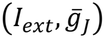 indicates that the SN structure is also generic over 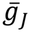 as a bifurcation parameter, suggesting additional multistability alongside the SN structure, either over *I*_*ext*_ or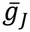. The observation of strongly depolarized front propagation in a 1-D array is an exciting new biophysical possibility and suggests spatio-temporal instabilities that produce voltage gradients and sustained LFPs within networks otherwise considered equipotential.

### Putting multistability and isopotentiality side by side

Isopotentiality is a state of the astrocytic network, and at the same time, its capability to restore that state after transient *V* changes, implying that any quasi-stable spatial *V* gradient would challenge the concept. Array simulations in Scenario 1 showed that, for *I*_*ext*_ and 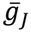 near the tip of the cusp (Fig. 5D), we observe *V*_*r*_ → *V*_*dr*_ front propagation. Since the analysis required bistable *f*, the more pronounced sensitivity of bistability to *g*_*J*_(*V*_*J*_) makes experimental validation of both effects particularly important. (i) Is it possible to design an experimental protocol to simulate any form of front propagation in an astrocytic preparation with deregulated Kir, while keeping the neurons quiet? (ii) How robust would the traveling depolarization be to small variations of 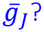 (iii) How does nonlinear *f* impact experimentally observed rapid restoration of isopotentiality, and does this impact change with 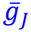 variation? Answers to these questions would cross-validate the model and experimental results, potentially qualifying *V* isopotentiality as a disease marker in pathologies characterized by strongly nonlinear steady-state *I*_*glia*_.

A provocative modeling question about the origin of multistability seems unavoidable. Could a variant of linear *f*, known to generate only a physiological *V*_*r*_ steady state in Eq. (6), end up being bistable only as an effect of nonlinear *g*_*J*_(*V*_*J*_)? If that is the case, it would provide a rationale for a promising combined, experimental and modeling study on physiologically healthy astrocytes. A two-electrode measurement setup would preliminarily inform the model about a realistic 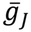 range. After producing the bistability, the model would inform the experiments about the *I*_*ext*_ needed to perturb *V*_*r*_ within its range and produce a *V*_*r*_ → *V*_*dr*_ transition to validate the restoration of isopotentiality. A variant of dynamic clamp protocol seems like an exciting new possibility for such a setup. Oscillating astrocytes Bifurcation scenarios for Scenario 2, with 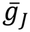 as a bifurcation parameter, demonstrate for the first time stable oscillatory behavior of membrane voltage in a glial model with *f* based on experimental data. The fact that robust SPO behavior could arise by increasing and decreasing 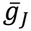 should not sound strange, given that *g*_*J*_(*V*_*j*_) inactivation is the main GJ gating mechanism and lowering 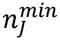 enabled the SPO structures in Scenario 2. A failure of front propagation in Fitzhugh–Nagumo arrays has also been demonstrated in the limit of weak coupling^54^.

There is strong experimental evidence from animal models that fully developed and prolonged epileptic seizures result in disconnection of hippocampal astrocytes^66^ in mice, implicating also changes in the expression of dominant Kir4.1 currents and real-time changes in gap–junctional properties^67^. In our view, the level of detail incorporated in Eq. (2) and Eq. (6) could motivate modeling such disconnection in temporal lobe epilepsy. Counterintuitively, molecular studies of mice genetically deficient in Cx43 and Cx30 connexins showed that the loss of those subtypes still leaves the astrocytes with intact cell-autonomous functions, such as glutamate and *K*^+^uptake, but impairs network integrity, as indicated by a reduction in short-term synaptic plasticity^68^.

The case of non-spiking astrocytes probably requires a slight change in how multistability is perceived. While the erratic bistable switching is commonly considered a transient instability^13^, the birth of stable spiking could also be perceived as a form of metastability, with *V* undergoing small-amplitude periodic changes in response to neural synchrony, with *I*_*ext*_ representing the impact of LFPs and [*K*^+^]_*o*_ variations. However, physiologically speaking, all electrogenic responses of astrocytes studied so far have a homeostatic nature^1^—one directed towards maintaining a near-isopotential network, assisting neurons in maintaining Nernstian equilibrium potentials. From that perspective, even slow, low-amplitude, sustained periodic oscillations in the 10 Hz range, if they persist, should be considered dramatic non-physiological behavior. Such oscillations would dramatically disturb all active ion transport by modulating ion concentrations inside and outside the cells, with a very high metabolic cost if prolonged and sustained. If validated experimentally, periodic oscillations in astrocytes could become a prime suspect in linking astrocytes to certain pathological manifestations in the brain.

The illustrated bifurcations over *I*_*ext*_ and 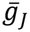 appeared to be stereotypical perturbations of the self-coupled cell. However, different, more atypical limit cycle behaviors are observed when varying other parameters, including saddle-node on invariant circle (SNIC) and subcritical HB bifurcations of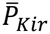, and saddle-homoclinic orbit (SHO) bifurcations of *ρ*_*r*_ and 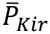, among others. They are structurally unstable boundaries in the parameter space around the central fold structure. See *Supplementary Material S6* for an example showing the coexistence of SNIC and SHO bifurcations at the boundaries of a fold domain and a supercritical Hopf bifurcation domain, within the same interval of *ρ*_*r*_.

### Slow changes and network plasticity

The plastic nature of gap junctions, with permeability changing in response to different phosphorylations^18,43^ of GJ proteins, further motivates biophysically novel model extensions. Exploring phosphorylation effects in Eq. (4) by numerical bifurcation analysis would potentially elucidate novel dependencies of GJ open-channel probability over slower timescales.

It remains a matter of open debate whether astrocytes form a single system-wide network (syncytium) or are organized into multiple networks (syncytia)^69^. Recent imaging studies^70^ support the existence of large, globally interconnected networks beyond localized cortical compartments^71^, in addition to the localized syncytia that have been the focus of most of quantitative studies so far.

The main objective of this modeling study was to further motivate innovative experiments to measure transient responses of coupled astrocytes, either by further developing real-time voltage imaging of glia^72^ or by combining soft electronics interfaces^73^ and the dynamic clamp for computationally controlled stimulations and readouts. Given the difficulties of recording transient responses in these cells, even without validated estimates of 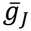in glia, we believe that we helped narrow the existing gap by introducing time- and voltage-dependent GJ junctions as a basis for future, more promising models.

### Model limitations

Even the simplest array model, a linear 1-D array of cells, is in principle not solvable, apart from trivial cases that are not of interest in biologically motivated studies. Analytical studies, reporting asymptotic analysis of simplified GJ-connected 1-D arrays^55^ illustrated that analyzing the first inner cell of the array should indicate the parameter ranges and nature of steady-state dependencies for observing jump transitions and front propagation. They also showed that opposite voltage gradients provide the key instabilities, with coupling strengths kept homogeneous over the array. This suggested that we could keep 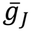 constant over the array in the initial model.

Even though we kept the model of self-coupled cell two-dimensional, both the complex cell kinetics *f* and the coupling term, taken together, present many model parameters, making the qualitative analysis of these ODEs over-defined in terms of parameters. For that reason, we stayed conservative with the parameter ranges of all measurable parameters, avoiding speculative conclusions that might arise if aberrant parameter values in numerical continuations suggested potentially interesting bifurcation structure.

## Supporting information

Supplementary Material - 7 sections

## Data availability

The *XPPAUT*^*57*^ .ode files and parameter files for the bifurcation analysis, as well as the MATLAB files for 1-D array simulations, will be available in *ModelDB* at https://senselab.med.yale.edu/ModelDB, under the model name “*Multistability in coupled astrocytes*”. The data dumps of particular bifurcation diagrams saved in AUTO / XPPAUT will also be available on request.

## Acknowledgements

This study was partly funded by the National Institute of Mental Health (US), grant number MH125030, and partly by the Association of National Science Organizations, grant number ANSO-CR-PP-2025-05.

## Footnotes

1 Numerically evaluating 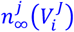 twice is only permissible in certain PDE numerical integration schemes where a perturbation at the left GJ precedes in time the change at the right GJ, which is by definition inactive at *t*_0_. In such a scheme, in addition to the main parallel integration step of the whole array, Δ*t*, a spatial propagation delay over the branched astrocyte structure should be defined, which is not a model assumption in our case.

2 Both packages, *AUTO/XPPAUT* and *MATCONT/MATLAB*, detected HB1, while only the first package extended the fold branches up to the limit point of cycles (LPC).

## References

1 A. Verkhratsky and M. Nedergaard, “Physiology of Astroglia,” Physiol Rev 98 (1), 239–389 (2018).

2 Alexei Verkhratsky and Arthur Butt, Glial neurobiology: a textbook. (John Wiley & Sons, 2007).

3 C. Giaume, A. Koulakoff, L. Roux, D. Holcman, and N. Rouach, “Astroglial networks: a step further in neuroglial and gliovascular interactions,” Nat Rev Neurosci 11 (2), 87–99 (2010).

4 M. Zhou, Y. Du, S. Aten, and D. Terman, “On the electrical passivity of astrocyte potassium conductance,” J Neurophysiol 126 (4), 1403–1419 (2021).

5 B. Ma, Buckalew, R., Du, Y., Kiyoshi, CM., Alford, CC., Wang, W., McTigue, DM., Enyeart JJ., Terman D., Zhou M., “Gap junction coupling confers isopotentiality on astrocyte syncytium,” Glia 64, 214–226 (2016).

6 S. Aten, C. M. Kiyoshi, E. P. Arzola, J. A. Patterson, A. T. Taylor, Y. Du, A. M. Guiher, M. Philip, E. G. Camacho, D. Mediratta, K. Collins, K. Boni, S. A. Garcia, R. Kumar, A. N. Drake, A. Hegazi, L. Trank, E. Benson, G. Kidd, D. Terman, and M. Zhou, “Ultrastructural view of astrocyte arborization, astrocyte-astrocyte and astrocyte-synapse contacts, intracellular vesicle-like structures, and mitochondrial network,” Prog Neurobiol 213, 102264 (2022).

7 G. E. Sosinsky and B. J. Nicholson, “Structural organization of gap junction channels,” Biochim Biophys Acta 1711 (2), 99–125 (2005).

8 G. Sosinsky, “Gap junction structure: New structures and new insights”, in Gap Junctions - Current topics in membranes (Elsevier, 2000), Vol. 49, pp. 1–22.

9 M. Simard and M. Nedergaard, “The neurobiology of glia in the context of water and ion homeostasis,” Neuroscience 129 (4), 877–896 (2004).

10 A G Pacholko, C A Wotton, and K K Bekar, “Astrocytes—the ultimate effectors of long-range neuromodulatory networks?” Frontiers in Cellular Neuroscience 14, 581075 (2020).

11 C. Steinhäuser, Seifert, G., Deitmer, J.W., “Physiology of astrocytes: ionchannels and ion transporters”, in Neuroglia (Oxford University Press, 2013), pp. 185–196.

12 G. Seifert, C. Henneberger, and C. Steinhäuser, “Diversity of astrocyte potassium channels - An update,” Brain Res Rev 126, 26–36 (2018).

13 P. Janjic, D. Solev, and L. Kocarev, “Non-trivial dynamics in a model of glial membrane voltage driven by open potassium pores,” Biophys J 122 (8), 1470–1490 (2023).

14 C. M. Kiyoshi, Y. Du, S. Zhong, W. Wang, A. T. Taylor, B. Xiong, B. Ma, D. Terman, and M. Zhou, “Syncytial isopotentiality: A system-wide electrical feature of astrocytic networks in the brain,” Glia 66 (12), 2756–2769 (2018).

15 S. Zhong, C. M. Kiyoshi, Y. Du, W. Wang, Y. Luo, X. Wu, A. T. Taylor, B. Ma, S. Aten, X. Liu, and M. Zhou, “Genesis of a functional astrocyte syncytium in the developing mouse hippocampus,” Glia 71 (4), 1081–1098 (2023).

16 S. Zhong, Y. Du, C. M. Kiyoshi, B. Ma, C. C. Alford, Q. Wang, Y. Yang, X. Liu, and M. Zhou, “Electrophysiological behavior of neonatal astrocytes in hippocampal stratum radiatum,” Mol Brain 9, 34 (2016).

17 G. P. Schools, M. Zhou, and H. K. Kimelberg, “Development of gap junctions in hippocampal astrocytes: evidence that whole cell electrophysiological phenotype is an intrinsic property of the individual cell,” J Neurophysiol 96 (3), 1383–1392 (2006).

18 D. A. Goodenough and D. L. Paul, “Gap junctions,” Cold Spring Harb Perspect Biol 1 (1), a002576 (2009).

19 M. Nuriya, A. Morita, T. Shinotsuka, T. Yamada, and M. Yasui, “Norepinephrine induces rapid and long-lasting phosphorylation and redistribution of connexin 43 in cortical astrocytes,” Biochem Biophys Res Commun 504 (4), 690–697 (2018).

20 S. Anders, D. Minge, S. Griemsmann, M. K. Herde, C. Steinhauser, and C. Henneberger, “Spatial properties of astrocyte gap junction coupling in the rat hippocampus,” Philos Trans R Soc Lond B Biol Sci 369 (1654), 20130600 (2014).

21 T. Manninen, J. Acimovic, and M. L. Linne, “Analysis of Network Models with Neuron-Astrocyte Interactions,” Neuroinformatics 21 (2), 375–406 (2023).

22 E. Dossi, L. Zonca, H. Pivonkova, G. Milior, J. Moulard, L. Vargova, O. Chever, D. Holcman, and N. Rouach, “Astroglial gap junctions strengthen hippocampal network activity by sustaining afterhyperpolarization via KCNQ channels,” Cell Rep 43 (5), 114158 (2024).

23 R D’Ambrosio, D S Gordon, and H. R. Winn, “Differential role of KIR channel and Na+/K+-pump in the regulation of extracellular K+ in rat hippocampus,” Journal of neurophysiology 87 (1), 87–102 (2002).

24 B S Nielsen, B R Larsen, A B Ghazal, A Katz, KC Brennan, S JD Karlish, and N. MacAulay, “Glial Versus Neuronal Na+/K+-ATPase in Activity-Evoked K+ Clearance and Their Sensitivity to Elevated Extracellular K+,” Glia (2025).

25 A Reichenbach, D Dettmer, W Reichelt, and W Eberhardt, “High Na+ affinity of the Na+, K+ pump in isolated rabbit retinal Müller (glial) cells,” Neuroscience letters 75 (2), 157–162 (1987).

26 G. Seifert, K. Hüttmann, D.K. Binder, C. Hartmann, A. Wyczynski, C. Neusch, and C. Steinhäuser, “Analysis of Astroglial K+ Channel Expression in the Developing Hippocampus Reveals a Predominant Role of the Kir4.1 Subunit,” J. Neurosci. 29, 7474–7488 (2009).

27 A. Destexhe, Huguenrad, J.,, “Nonlinear Thermodynamic Models of Voltage-Dependent Currents,” J Comp Neurosci 9, 259–270 (2000).

28 M.B. Jackson, Molecular and Cellular Biophysics. (Cambridge Univ. Press, New York, 2006).

29 R. H. Adrian, “Rectification in muscle membrane,” Prog Biophys Mol Biol 19 (2), 339–369 (1969).

30 D. Zuo, Chen, K., Zhou, M., Liu, Z., Chen, H, “Kir2.1 and K2P1 channels reconstitute two levels of resting membrane potential in cardiomyocytes,” J Physiol 595, 5129–5142 (2017).

31 H. Chen, Zuo, D., Zhang, J., Zhou, M., Ma, L., “Classification of 2-pore domain potassium channels based on rectification under quasi-physiological ionic conditions,” Channels 8, 503–508 (2014).

32 R. Wilders and H. J. Jongsma, “Limitations of the dual voltage clamp method in assaying conductance and kinetics of gap junction channels,” Biophys J 63 (4), 942–953 (1992).

33 M. V. Bennett and V. K. Verselis, “Biophysics of gap junctions,” Semin Cell Biol 3 (1), 29–47 (1992).

34 R. Vogel, V. Valiunas, and R. Weingart, “Subconductance states of Cx30 gap junction channels: data from transfected HeLa cells versus data from a mathematical model,” Biophys J 91 (6), 2337–2348 (2006).

35 R. D. Veenstra, “Comparative physiology of cardiac gap junction channels”, in Biophysics of Gap Junction Channels (CRC Press, 2018), pp. 131–144.

36 J. I. Nagy and J. E. Rash, “Connexins and gap junctions of astrocytes and oligodendrocytes in the CNS,” Brain Res Brain Res Rev 32 (1), 29–44 (2000).

37 R. Vogel and R. Weingart, “The electrophysiology of gap junctions and gap junction channels and their mathematical modelling,” Biol Cell 94 (7-8), 501–510 (2002).

38 T.A. Bargiello, S. Oh, Q. Tang, N.K. Bargiello, T.L. Dowd, and T. Kwon, “Gating of Connexin Channels by transjunctional-voltage: Conformations and models of open and closed states,” Biochim. Biophys. Acta 1860, 22–39 (2018).

39 S. Oh, and Bargiello, T.,, “Voltage Regulation of Connexin Channel Conductance,” Yonsei Med J 56, 1–15 (2015).

40 C. Peracchia, “Gap Junction Channel Regulation: A Tale of Two Gates-Voltage Sensitivity of the Chemical Gate and Chemical Sensitivity of the Fast Voltage Gate,” Int J Mol Sci 25 (2) (2024).

41 H. Jongsma, R. Wilders, A.C.G. van Ginneken, and M.B. Rock, “Modulatory effect of the transcellular electrical field on gap junctional conductance”, in Biophysics of Gap Junction Channels, edited by C. Peracchia (CRC Press, Boca Raton, FL, 1991), pp. 163–172.

42 L. C. Barrio, T. Suchyna, T. Bargiello, L. X. Xu, R. S. Roginski, M. V. Bennett, and B. J. Nicholson, “Gap junctions formed by connexins 26 and 32 alone and in combination are differently affected by applied voltage,” Proc Natl Acad Sci U S A 88 (19), 8410–8414 (1991).

43 A. P. Moreno, G. I. Fishman, and D. C. Spray, “Phosphorylation shifts unitary conductance and modifies voltage dependent kinetics of human connexin43 gap junction channels,” Biophys J 62 (1), 51–53 (1992).

44 S. Baigent, J. Stark, and A. Warner, “Convergent dynamics of two cells coupled by a nonlinear gap junction,” J Nonlinear Analysis 47 (1), 257–268 (2001).

45 A. L. Harris, D. C. Spray, and M. V. Bennett, “Kinetic properties of a voltage-dependent junctional conductance,” J Gen Physiol 77 (1), 95–117 (1981).

46 A. L. Harris, D. C. Spray, and M. V. Bennett, “Control of intercellular communication by voltage dependence of gap junctional conductance,” J Neurosci 3 (1), 79–100 (1983); D. C. Spray, A. L. Harris, and M. V. Bennett, “Equilibrium properties of a voltage-dependent junctional conductance,” J Gen Physiol 77 (1), 77–93 (1981).

47 R. Vogel and R. Weingart, “Mathematical model of vertebrate gap junctions derived from electrical measurements on homotypic and heterotypic channels,” J Physiol 510 (Pt 1) (Pt 1), 177–189 (1998).

48 V. K. Verselis, M. V. Bennett, and T. A. Bargiello, “A voltage-dependent gap junction in Drosophila melanogaster,” Biophys J 59 (1), 114–126 (1991).

49 N. A. Oberheim, S. A. Goldman, and M. Nedergaard, “Heterogeneity of astrocytic form and function,” Methods Mol Biol 814, 23–45 (2012).

50 E. A. Bushong, M. E. Martone, Y. Z. Jones, and M. H. Ellisman, “Protoplasmic astrocytes in CA1 stratum radiatum occupy separate anatomical domains,” J Neurosci 22 (1), 183–192 (2002).

51 James P. Keener and James Sneyd, “Mathematical physiology”, in Interdisciplinary applied mathematics v 8 (Springer,, New York, 1998).

52 J. D. Murray, “Mathematical biology”, in Biomathematics v 19 (Springer-Verlag,, Berlin ; New York, 1993).

53 K. Kladko, I. Mitkov, and A.R. Bishop, “Universal scaling of wave propagation failure in arrays of coupled nonlinear cells,” Phys Rev Lett 84 (19), 4505 (2000).

54 T. Erneux and G. Nicolis, “Propagating waves in discrete bistable reaction-diffusion systems,” Physica D 67 (1-3), 237–244 (1993).

55 V. Booth and T. Erneux, “Understanding propagation failure as a slow capture near a limit point,” SIAM J Appl Math 55 (5), 1372–1389 (1995).

56 I. Rajapakse and S. Smale, “Emergence of function from coordinated cells in a tissue,” Proc Natl Acad Sci U S A 114 (7), 1462–1467 (2017).

57 B. Ermentrout, Simulating, analyzing, and animating dynamical systems: a guide to XPPAUT for researchers and students. (Society of Industrial and Applied Mathematics (SIAM), Philadelphia, 2002).

58 E. Doedel, A.R. Champneys, T.F. Fairgrieve, Y.A. Kuznetsov, B. Sandstede, and X. Wang, AUTO 97: Continuation and bifurcation software for ordinary differential equations (with HomCont) (indy.cs.concordia.ca, 1997).

59 A. Dhooge, W. Govaerts, and Yu A Kuznetsov, “MATCONT: a MATLAB package for numerical bifurcation analysis of ODEs,” ACM Transactions on Mathematical Software (TOMS) 29 (2), 141–164 (2003).

60 E.M. Izhikevich, Dynamical Systems in Neuroscience - The Geometry of Excitability and Bursting. (The MIT Press, Cambridge, Massachusetts, 2007).

61 S. E. Nwaobi, V. A. Cuddapah, K. C. Patterson, A. C. Randolph, and M. L. Olsen, “The role of glial-specific Kir4.1 in normal and pathological states of the CNS,” Acta Neuropathol 132 (1), 1–21 (2016).

62 Y. Ohno, N. Kunisawa, and S. Shimizu, “Emerging Roles of Astrocyte Kir4.1 Channels in the Pathogenesis and Treatment of Brain Diseases,” Int J Mol Sci 22 (19) (2021).

63 K Bogus, Nicoletta Marchesi, L IM Campagnoli, A Pascale, and A Pałasz, “Glial Cells as Key Mediators in the Pathophysiology of Neurodegenerative Diseases,” International Journal of Molecular Sciences 27 (2), 884 (2026).

64 M. Goldberg, M. De Pitta, V. Volman, H. Berry, and E. Ben-Jacob, “Nonlinear gap junctions enable long-distance propagation of pulsating calcium waves in astrocyte networks,” PLoS Comput Biol 6 (8) (2010).

65 V. Matrosov, S. Gordleeva, N. Boldyreva, E. Ben-Jacob, V. Kazantsev, and M. De Pittà, “Emergence of regular and complex calcium oscillations by inositol 1, 4, 5-trisphosphate signaling in astrocytes”, in Computational glioscience (Springer, 2019), pp. 151–176.

66 P. Bedner, A. Dupper, K. Huttmann, J. Muller, M.K. Herde, P. Dublin, T. Deshpande, J. Schramm, U. Haussler, C.A. Haas, C. Henneberger, M. Theis, and C. Steinhäuser, “Astrocyte uncoupling as a cause of human temporal lobe epilepsy,” Brain 138, 1208–1222 (2015); Seifert G. and Bedner P. Steinhäuser C., “Astrocyte Dysfunction in Temporal Lobe Epilepsy: K+ Channels and Gap Junction Coupling,” GLIA, 60:1192 (2012).

67 T. Deshpande, T. Li, M. K. Herde, A. Becker, H. Vatter, M. K. Schwarz, C. Henneberger, C. Steinhauser, and P. Bedner, “Subcellular reorganization and altered phosphorylation of the astrocytic gap junction protein connexin43 in human and experimental temporal lobe epilepsy,” Glia 65 (11), 1809–1820 (2017).

68 U. Pannasch, L. Vargova, J. Reingruber, P. Ezan, D. Holcman, C. Giaume, E. Sykova, and N. Rouach, “Astroglial networks scale synaptic activity and plasticity,” Proc Natl Acad Sci U S A 108 (20), 8467–8472 (2011).

69 V. Houades, A. Koulakoff, P. Ezan, I. Seif, and C. Giaume, “Gap junction-mediated astrocytic networks in the mouse barrel cortex,” J Neurosci 28 (20), 5207–5217 (2008).

70 M. L. Cooper, M. C. Selles, M. Cammer, C. Redd, H. K. Gildea, J. Sall, K. E. Chiurri, P. Cheung, D. G. Wheeler, A. S. Saab, S. A. Liddelow, and M. V. Chao, “Astrocytes connect specific brain regions through plastic networks,” Nature (2026).

71 J. Visser, G. Milior, R. Breton, J. Moulard, M. Garnero, P. Ezan, J. Ribot, and N. Rouach, “Astroglial networks control visual responses of superior collicular neurons and sensory-motor behavior,” Cell Rep 43 (7), 114504 (2024).

72 M. Armbruster, S. Naskar, J. P. Garcia, M. Sommer, E. Kim, Y. Adam, P. G. Haydon, E. S. Boyden, A. E. Cohen, and C. G. Dulla, “Neuronal activity drives pathway-specific depolarization of peripheral astrocyte processes,” Nat Neurosci 25 (5), 607–616 (2022).

73 L. Maiolo, V. Guarino, E. Saracino, A. Convertino, M. Melucci, M. Muccini, L. Ambrosio, R. Zamboni, and V. Benfenati, “Glial Interfaces: Advanced Materials and Devices to Uncover the Role of Astroglial Cells in Brain Function and Dysfunction,” Adv Health Mater 10 (1), e2001268 (2021).

