## Supplementary Material - 7 sections for "Membrane voltage multistability in coupled glial cells"

to accompany

### Supplementary Material S1: Gap junctions and gap junction channels

Gap junctions (GJs) are formed by clusters of hundreds to approximately two thousand gap junction channels (GJCs) in a single membrane cluster[1] or a plaque (Fig. S1C). Each plaque represents one of multiple connections between two cells, or, electrically, one high-conductance junction. A single conductive GJ channel (a single GJC) between the two glial cells is formed when a pair of channel units (connexons), one at each side, dock (axially aligned; Fig. S1B) to form a single conductive pore/path. Each connexon may have more than one conductive state, giving rise to complex gating transitions in some variants. *Connexins* (abbreviated Cx) are the family of proteins that build the channel pore on each side. Figure S1B shows the elements of this structural hierarchy. They form either homotypic GJCs, in which both channels are made by the same connexin subtype, or heterotypic channels assembled from two different connexins, meaning different GJCs have different conductive properties.

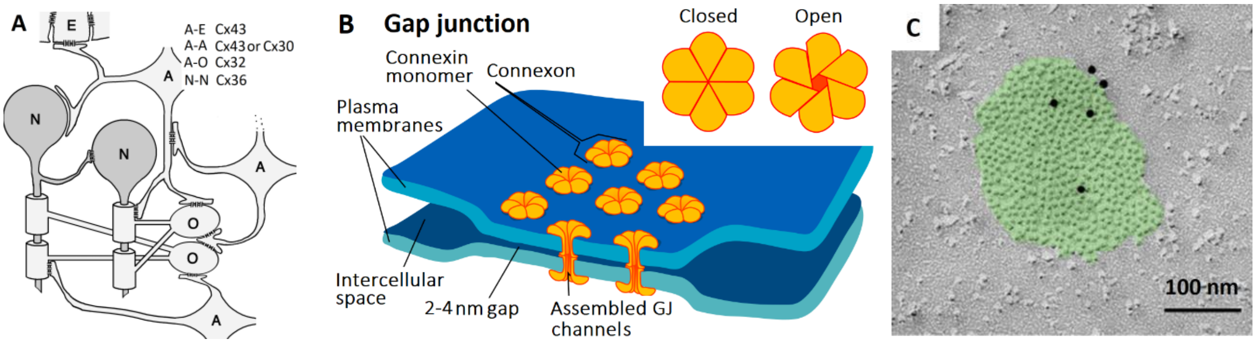


**Figure S1. Gap junctions (GJ) and gap junction channels (GJCs). (A)** Depending on the connection, astrocytes assemble different GJC combinations. Astrocytes dominate the glial networks and are connected by homotypic Cx43–Cx43 or Cx30–Cx30 channels, but they also connect directly to other glial cell types, typically oligodendrocytes. The cell designations are as follows: "A" an astrocyte; "O" an oligodendrocyte; "N" a neuron; and "E" an endothelial cell of the blood vessels or an ependymal cell at the external boundaries of the. Image from Rash J. et al (2001)[2], reproduced with permission from the *Society for Neuroscience*. **(B)** Hierarchical structure of the macroscopic gap junction, starting with Connexin protein units forming a single channel, and extending to a physical cluster of docked channels forming one junction among many. An open GJC pore is $\sim 2nm$ wide, which is 5–10 times wider than the pores in most voltage-gated ion channels. The drawing is adapted from Wikipedia, illustrated by Mariana Ruiz Villarreal. **(C)** Freeze-fraction electron micrograph of a plaque of channels forming a single gap junction connection, as seen on the cell surface of one of the connecting astrocytes (the other having been experimentally detached). Black dots are gold particles that appear as an experimental artefact of the labeling method. Such a structure may contain up to a few thousand channel pores. Image from J.C. Recktenwald, PhD thesis, 2020, University of Saarland, Homburg/Saar, Germany.

Sizes and channel densities within a single glial gap junction[3] suggest biophysical complexity that cannot be conceptualized with a single channel pore (Fig. S1C). For more structure-function details on GJ connections, see Sosinsky G. (2000)[4].

### Supplementary Material S2: **Equivalent circuit of recording setup from two GJ-connected cells**

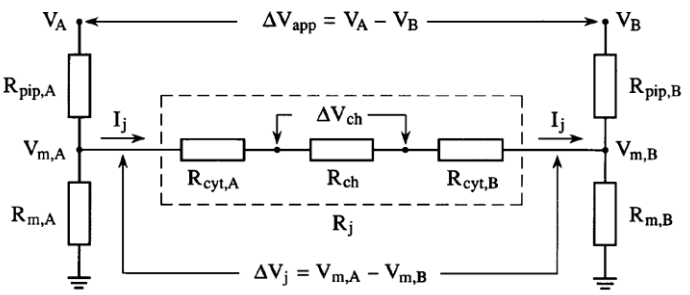


**Figure S2. Equivalent resistive circuit of a two-electrode whole-cell recording setup.** Each electrode, at $V_A$ and $V_B$, introduces a series pipette resistance $R_{pip}$ and a branching point for the current path through the GJ. Apart from the cell input resistance $R_m$, the passive properties of the junction define a cytoplasmic access resistance $R_{cyt}$, in addition to $R_{ch}$, the total ohmic resistance of the cluster of GJ channels across *all junctions*. These resistances are represented by a single resistor $R_j$, since $R_{cyt}$ and $R_{ch}$ are electrically inseparable. Designations are adopted from the circuit in Ref. [5]

In the equivalent circuit diagram, which is generic to any two-electrode setup on two directly connected cells[5] (Fig. S2), each electrode A and B measures the voltage drop over the divider $R_{pip} + R_m$, typical of a whole-cell voltage clamp. In addition, the net current through the gap junction(s) $I_j$, proportional to $V_{A,B}R_m \,/\, (R_{pip} + R_m\,)$ is subject to junction-specific cytoplasmic access resistance $R_{cyt}$ in addition to $R_{ch}$, the total resistance of the plaque as a two-terminal ohmic element. The $R_{cyt}$ resistance physically does not simply represent a fraction of the whole-cell input resistance $R_m$, but is a specifically defined resistance arising from the resistive properties of the junctional plaque as a “connector device”[5] when fully conductive. $R_{cyt}$ is not known, since it is impossible to measure or sum it for all junctions, due to its size dependence[6]. Yet, it effectively introduces fractional effects of $R_{cyt}/R_m$ on $V_A$, and $V_B$, or on the *apparent voltage* difference $\Delta V_{app} = V_A - V_B$ that polarizes the junction(s). For more details on both the static and kinetic effects of the uncertainty in $R_{cyt}$ and $R_{ch}$, see Wilders and Jongsma, 1992[5]. Because it is impossible to distinguish experimentally between the contributions of $R_{cyt}$ and $R_{ch}$, we use $R_j$ to represent the resistance of an entire junction, as a “symmetrical” resistance seen from either of the cells. Apart from the artifactual resistances, we define the actual *transjunctional voltage* as $V_j = V_{m,A} - V_{m,B}$, by convention taken as positive when $(V_{m,OTEHR} - V_{m,OWN})$ is positive. In the rest of our model development, we will also move from apparent resistance to conductances, as is conventional in conductance-based descriptions of neural circuits.

Despite the single-sided $V_m$ dependence of the connexons—GJC half-channels (Fig. S1 has demonstrated that $V_m$ gates each of them independently)[7]—their effects on $V_J$ gating could not be studied in a native setup with a fully functional GJ, but are known to produce a minor effect on principal $V_J$ gating[8,9]. In the model development in the main text, we describe only $V_J$-dependences.

### Supplementary Material S3: **Adjusting maximal Kir permeability in an *effective cell cluster***

Figure S3 illustrates the effect of an increase in Kir channel maximal Nernst-Planck permeability $\bar{P}_{Kir}$ on the shape of total glial current $I_{glia}$ , in case of pronounced $I_{Kir}$ rectification, $\Delta z_B < -1$ (see the description of Kir current[10] or Eq. (2) in the main text). In this way, we get closer to the actually measured $I_{Kir}$ currents and maximal slope conductances, $\bar{g}^c_{Kir}$, in an intact tissue: a voltage-clamped cell with a fraction of active GJs connecting branches of immediately neighboring cells, thereby defining an effectively measured cell cluster.

The $I_{Kir}$ profile (Figure S3A, blue; the same as Fig. 2C in the main text) displays a close to 1.5-fold increase in $\bar{P}_{Kir}$compared with the value obtained in the model of an isolated cell[10], after adjustment for $[K^+]_o$ dependence; see Section 7.1 of *Supplementary Information 7*. In Fig. S3B, a family of $I_{Kir}$ curves is shown for $\bar{P}_{Kir}$ over a wide range, between 1e-07 and 9e-07 $cm^3$/s, where the upper range should describe the maximal channel densities in the glial end-feet or the thin processes enwrapping the synapses. We chose this range to reflect the real channel densities suggested by voltage-imaging and expression studies[11], rather than using it as a numerical stretch to construct N-shaped curves.

As a parameter sanity check, approximating the equivalent maximal slope conductance from the close-to-linear part of the curve, between -110 and -60 mV, gave a value close to **50 nS**, which is well within the range of $\bar{g}^c_{Kir}$ values estimated from single-cell *in situ* recordings with a cell capacitance in the 20–50 pF range across multiple studies.

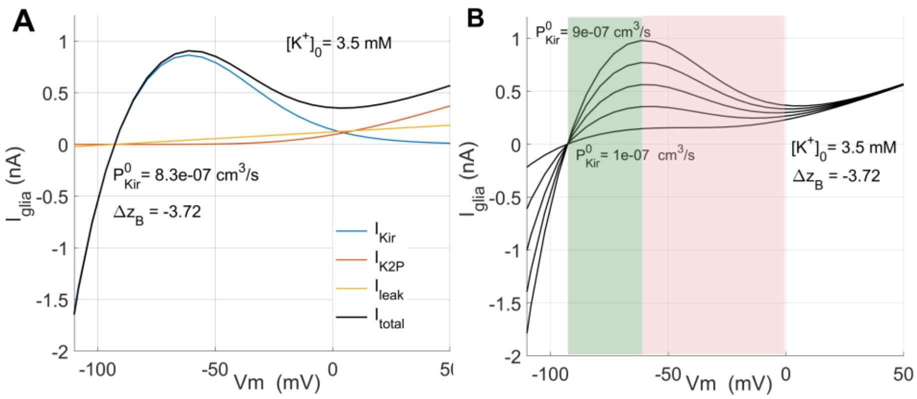


**Figure S3. I–V profiles of $\boldsymbol{I_{glia}}$for larger $\boldsymbol{P_{Kir}}$.** (A) Example of a steady state I–V profile of glial current, with a 1.5-fold increase in $\bar{P}_{Kir}$ in the Nernst-Planck description, (Eq. (2)), compared with that of an isolated cell (Janjic et al., 2023)[10], with $\Delta z_B = -3.72$. (B) Family of I–V curves for $\bar{P}_{Kir}$in the range 1e-07 to 9e-07 $cm^3$/s. The red-marked region shows the negative slope, where $I_{kir}$ develops an N-shape.

This illustrates that in addition to the dominant effect of $\Delta z_B$ on $I_{Kir}$ nonlinearity, increasing $\bar{P}_{Kir}$also pronounces the nonlinearity in the same outward range. Either change in $I_{Kir}$ nonlinearity in Fig. S3 produces steeper negative slopes in the I–V curve and steeper linear segments within the $V_m$ range below $V_r$, both making the depolarization-induced transitions more likely. In other words, smaller depolarizations will move the state from the positive I–V slope (the nominal, green-shaded WR profile) over the hump to the negative-slope transitory range (red-shaded range), knocking the cell onto the upper branch of the fold curve and into the depolarized state $V_{dr}$. Even though they are not presented in detail, enriched bifurcation scenarios around the fold structure, as an organizing center, were observed under multiple parameter variations, including $\bar{P}_{Kir}$: see comments in *Supplementary Information S5.*

Supplementary Material S4: **Nonlinear Least Squares curve fitting of UPO frequency dependence on $\bar{\boldsymbol{g}}_{\boldsymbol{j}}$ in Case 2 bifurcation scenario**

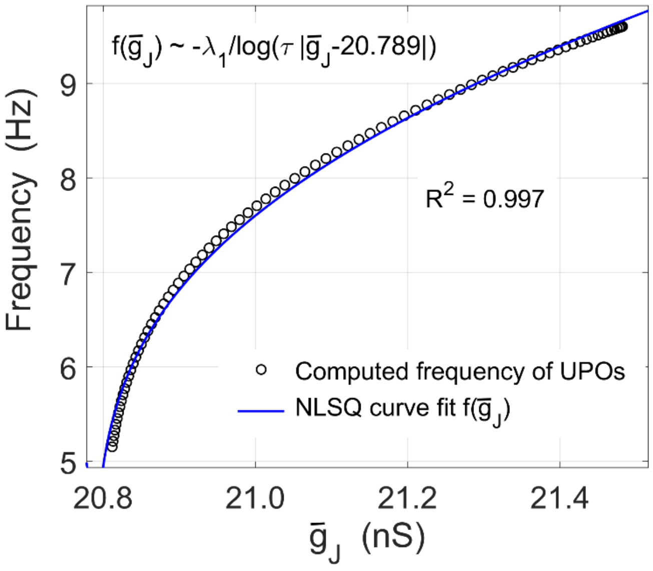


**Figure S4. UPO Frequency dependence on $\bar{\boldsymbol{g}}_{\boldsymbol{gj}}$.** The UPO curve approaches zero as $\lambda_1 / \log\left(\tau * \left|\bar{g}_J - \bar{g}_J^{LPC}\right|\right)$ as $\bar{g}_J \rightarrow \bar{g}_J^{LPC} = 20.789\ nS$, representing the LPC value of the bifurcation parameter $\bar{g}_{gj}$ in Fig. 7A and 7B. This curve is known to be a feature of the saddle homoclinic orbit bifurcation (SHO)[12] scenario, which we did not detect numerically. For the positive eigenvalue $\boldsymbol{\lambda_1} = \mathbf{0.04242}$ of the saddle and the negative saddle quantity $(\lambda_1 + \lambda_2)$, it corresponds to a band of UPOs. This suggests a possible origin of the bifurcation picture in Case 2, Fig. 7A, where the fold and UPO structures could be attractor“ruins“ of an SHO structure collapsing when it meets either the SPO or the fold structure. $\tau$=17.92, obtained by fitting, is a multiplier depending on the size of the domain of the phase plane containing the UPO, as well as other global features of $f$[12].

Supplementary Material S5: **Small-amplitude *SPO* bifurcation over $\bar{g}_J$.**

Changing the slope factor of $n_J$ to $v_S = 7\ mV$, within the $(\bar{g}_J, V_m)$ domain of the Case 2 bifurcation diagram introduced a small-orbit SPO, with a maximal amplitude of near 10 mV, again through a supercritical HB. Similar to Case 2, decreasing $\bar{g}_J$ through $HB_{right}$ or $\bar{g}_{gj} = 23.17\ nS$ , stable oscillations emerged. $V_r$ continuation along the same branch from the left, $HB_{left}$ or $\bar{g}_J = 22.15175\ nS$ detected a *fold limit cycle bifurcation*, Fig. S5A, with the blue-shaded region enlarged in Fig. S5B.

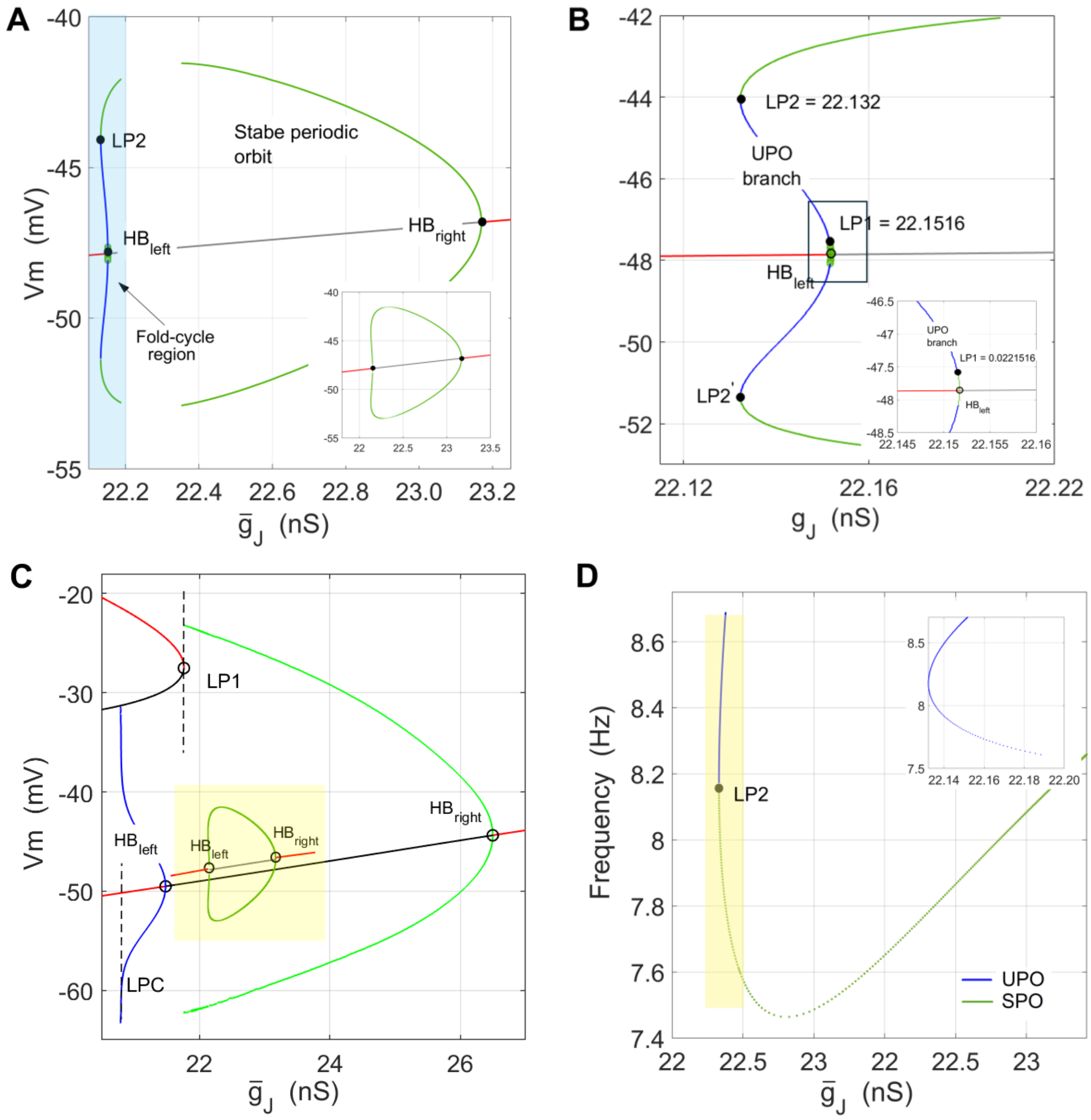


**Figure S5. Back-to-back limit cycle bifurcation. (A)** Bifurcation diagram $(\bar{g}_J, V_m)$ showing a region of stable limit cycles (SPOs) about 1 nS wide, emerging through (i) a supercritical HB at the $HB_{right}$ point, and disappearing via (ii) a *fold cycle bifurcation* at the $HB_{left}$ point. Running both continuations separately produced a closed loop; see inset. **(B)** The blue-shaded region in (A), 0.1 nS wide, is expanded, showing that in the fold cycle bifurcation, the radius of the orbit undergoes a fold bifurcation[12]. **(C)** The small-orbit SPO, indicated by the yellow-shaded square, shown within the bifurcation picture of the large-orbit SPO scenario of Case 1, Fig. 7A. **(D)** Frequency plot showing that the small-amplitude SPO in (A) has a non-zero onset frequency, in accordance with both transitions. Relevant changeable parameters are as follows: $V_{dr}^{l} = -20\ mV,\ V_r^r = -55.4\ mV,\ I_{ext} = 0.55\ nA, \bar{P}_{Kir} = 8.28 \times 10^{-7} cm^3/s\ ,\ V_{12-Kir} = -55\ mV, z_{BA} = -3.72,\ v_s = 7,\ [K^+]_o = 3.5\ mM,\ n_J^{min} = 0.25,\ n_J^{max} = 0.95,\ \rho_l = 0.8, \rho_r = 0.2.$

Even though this mechanistic description is not wrong in describing the actual continuation, the fold cycle orbit (FCO) bifurcation, also called a *dual limit cycle*, is a limit cycle bifurcation in which the SPO of the supercritical HB disappears. It is a very thin region of $0.02\ nS$, enlarged in Fig. S5B, in which the SPO collapses, and this required decreasing the continuation step in AUTO/XPPAUT to 0.005, from 0.01, which was used in obtaining the SPO branches from $HB_{right}$.

In Figure S5C, the yellow rectangle, shows this small-orbit SPO structure within the bifurcation diagram of the Case 1 SPO example. With its much smaller amplitude and energy cost, this SPO is a more plausible instability to be observed, or to remain undetected, than the large-orbit SPO in Case 2.

The continuity of both frequency arcs through LP2 in Fig. S5D, and the two symmetric amplitude arcs folding from $HB_{left}$ towards the limit points LP2 and LP2', show that both bifurcation structures belong to the same limit cycle structure[12]. The amplitude arcs folding through LP2 and LP2' illustrate that, through a fold cycle bifurcation, or saddle-node of limit cycles[12], the SPO amplitude undergoes a saddle-node bifurcation before disappearing into stable $V_r$. The non-zero terminating frequency of the UPOs, the blue curve in Fig. 9D and its inset, distinguishes this fold cycle bifurcation from a subcritical Hopf bifurcation.

### Supplementary Material S6: **Coexistence of SNIC bifurcation of a fixed point and saddle homoclinic orbit bifurcation of stable periodic orbit.**

Figure S7 shows an example of a rich one-parameter bifurcation structure over the range of $\rho_r$, the fraction parameter describing the stochastic variability of astrocytic GJ that does not meet the initially depolarized front in the 1-D array model, Eq. (6). This is one of multiple configurations around the fold structure (existing in $f$), introduced by the parameters of the coupling term.

In this example, instead of the band of UPOs as in Scenario 1 and Scenario 2, a parabolic domain outlined by the Hopf bifurcation curve has been nested along the stable branch of the fold structure, extending the unstable segment of the fold curve of $V_r$. This arrangement produces two tangencies near LP2 at $\rho_b = 111658$ organized by the higher-order codimension-2 bifurcation structure: (a) the SNIC as limit behavior over the fold structure, and (b) the saddle-homoclinic orbit bifurcation (SHO or HCO) of the SPO extending to the left of the Hopf bifurcation point. We say "near" because the corresponding branches are properly topologically organized in $\mathbb{R}^1 \times \mathbb{C}^2$ by Bogdanov–Takens and Bautin bifurcation structures. There is no proper boundary between these two structures in the $\mathbb{R}^1 \times \mathbb{C}^1$ plane. Zhou and Gu have described extensively these arrangements in Morris–Lecar model of neuronal autapse connection[13].

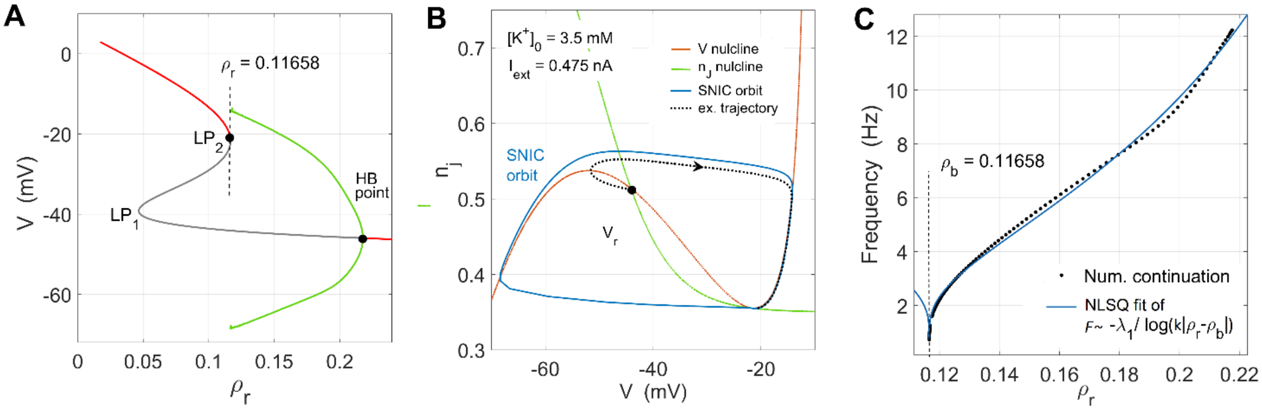


**Figure S7. Saddle-node on invariant circle and homoclinic orbit bifurcation**. **(A)** Bifurcation diagram over $\rho_r$ displaying coexistence of a fold and Hopf bifurcation region, meeting near the saddle at $-20\ mV$ and $\rho_r = 0.11658$, labeled LP2. Decreasing $\rho_r$ close to zero, which has a decoupling effect in Eq. (6) like decreasing $\bar{g}_J$ takes the state to the depolarized $V_{dr}$ as a sole steady-state, represented by the red branch. **(B)** The SNIC curve appearing at LP2, with the vanishing eigenvalue $\lambda_1$ near the tangency estimated in the range of $(0.0095 - 0.013)$. An example of a phase-plane trajectory is shown to illustrate the limit-cycle behavior of the SNIC. **(C)** Frequency dependence near the boundary when decreasing $\rho_r$, illustrating the limit-cycle bifurcation in which the SPO disappears through a homoclinic orbit bifurcation. Relevant changeable parameters are as follows: $\bar{g}_J = 18.6\ nS, \tau_j = 100\ ms, V_{dr}^l = -10\ mV,\ V_r^r = -55\ mV,\ I_{ext} = 0.475\ nA, \bar{P}_{Kir} = 9.0 \times 10^{-7} cm^3/s\ ,\ V_{12-Kir} = -55\ mV, z_{BA} = -3.68,\ v_s = 6,\ [K^+]_o = 3.5\ mM,\ n_J^{min} = 0.35,\ n_J^{max} = 0.95,\ \rho_l = 0.8.$

Supplementary Material S7: **Model equations and parameter tables**

### S7.1 Equations of glial currents in Eq. (3), main text

The values of parameters varied within the bifurcation analysis and simulations are explicitly given within Chapter 3.

**$I_{Kir}$ current** (parameters defined in the main text)

$$I_{Kir} = zFP_{Kir}\left([K^+]_i - [K^+]_o \exp\left(-\frac{dz\,V_m}{v_T}\right)\right) \Big/ \left(exp\left(\frac{-\delta(\Delta z_B + dz)(V_m - V_{12-Kir})}{v_T}\right) + 1\right) \quad (nA)$$

$$P_{Kir}(K_o) = \bar{P}_{Kir}(1 + S_{Kir}\, ln([K^+]_o/[K^+]_i),$$

The full model of $I_{Kir}$[10] allows variable permeability $P_{Kir}$ with $[K^+]_o$. Since we keep ion concentration constant in simulations and bifurcation analysis, we only list parameters of this dependence here. We refer interested readers to the full model for $[K^+]_o$-dependence for more details [10].

| Parameter | Units | Description | Value, default | Comment |
|---|---|---|---|---|
| $T$ | $K$ | Temperature | 298 | Experimental room temperature, 25˚C in Ref.[14] |
| $F$ | $C/mol$ | Faraday constant | 96485 | |
| $R$ | $J/mol \cdot K$ | Universal gas constant | 8.314 | |
| $v_T$ | $mV$ | Slope factor, $RT/F$ | 25.7 | at $T = 298°K$, constant in whole study |
| $[K^+]_o$ | $mM$ | External $K^+$concentration | 3.5 | |
| $[K^+]_i$ | $mM$ | Astrocytic, internal $K^+$concentration | 130 | |
| $C_m$ | $pF$ | Astrocytic membrane capacitance | 20 | Mean effective capacitance of the patch, whole-cell |
| $z$ | | Unitary valence of permeant $K^+$ion | 1.0 | |
| $dz$ | | Fractional valence of $K^+$ions | 1.0 | May have non-unitary value |
| $\bar{P}_{Kir}$ | $cm^3/s$ | Maximal permeability of $Kir$ channel | 8.28e-07 | Value used in Case-1 and 2. Varied in bifurcation analysis |
| $S_{Kir}$ | | Scaling parameter in $P_{Kir}([K^+]_o)$ | 0.23 | Used in fitting $[K^+]_o$ dependence of $P_{Kir}$ |
| $V_{12-Kir}$ | $mV$ | Half-activation voltage of $I_{Kir}$ | -55 | Boltzmann term in $p_{Kir}$ , Eq. 3 |
| $\Delta z_B$ | | Eq. charge of the cation block | -3.72 | Effective valence of blocking cation(s) |
| $\delta$ | | Fractional electrical length of $Kir$ pore | 0.5 | Describes the fraction of voltage drop of $V_m$ |

**$I_{K2P}$ current**

$$I_{K2P} = n_{K2P}^k\, I_{K2p-GHK}$$

$$= n_{K2P}^k\, P_{K2P}\frac{F^2 z_{K2P}^2}{RT} V_m \frac{([K^+]_i - [K^+]_o \exp(-z_{K2P}V_m/v_s))}{(1-\exp(-z_{K2P}V_m/v_s))}\ (nA).$$

| | | | | |
|---|---|---|---|---|
| $V_{12-K2p}^0$ | $mV$ | Half activation voltage $V_{12}$ at $[K^+]_o^{nom}$ | -20.5 | See $n_{K2P}$ activation. Not used in reduced model, Eq. (6) |
| $k$ | | k-th power in K2P activation kinetics | 2 | See $n_{K2P}$ activation. Not used in reduced model, Eq. (6) |
| $z_{K2P}$ | | Charge valence for K2P pores, fixed | 1.0 | $I_{K2P}$ model, GHK description, |
| $\tau_{K2P}$ | $ms$ | Activation time of K2P-TREK pore | 3.0 | See $n_{K2P}$ activation. Not used in reduced model, Eq. (6) |
| $P_{K2P}^b$ | $cm/s$ | K2P channel permeability, basal | 1.44e-08 | $I_{K2P}$ model, GHK description, |

#### $I_{leak}$, unspecific $K^+$ leak current

| | | | | |
|---|---|---|---|---|
| $g_{leak}$ | $nS$ | Glial leak conductance, Ohmic | 1.28 | Non-specific $K^+$ leak, |
| $E_K$ | $mV$ | Nernst reversal potential for $K^+$ | -92.9 | For $K^+$concentrations given in $I_{Kir}$ table |

### S7.2 Self-coupled cell: reduced model, Eq. (6) and GJ inactivation Eq. (4)

$$C_m dV_m/dt = f(V_m, I_{ext}) + \rho_l \, \bar{g}_J n_J (V_{dr} - V_m) + \rho_r \, \bar{g}_J n^J_{max} (V_r - V_m), \qquad (nA), \qquad (6a)$$

$$dn_J/dt = \left(n^J_\infty(V_J) - n_J\right)/\tau_J, \qquad (6b)$$

$$n^J_\infty(V_j) \equiv g_J(V_J)/\bar{g}_j = n^{min}_J + (n^{max}_J - n^{min}_J)/(1 + \exp(-(V_J - V_{j12})/v_S)), \qquad (4b)$$

| Parameter | Units | Description | Value (default) | Comment |
|---|---|---|---|---|
| $\bar{g}_j$ | $nS$ | Maximal GJ slope conductance | 18 to 27 | Bifurcation parameter |
| $n^{min}_J$ | | Min., residual GJ inactivation probability $n_J$ | 0.25 to 0.35 | Varied within bifurcation analysis |
| $n^{max}_J$ | | Max. GJ inactivation probability $n_J$ | 0.95 | |
| $V_{j12}$ | $mV$ | Half-activation $V_J$ voltage in GJ inactivation $n_J$ | 40 | See Chapter 2.1 for definition. |
| $v_S$ | $mV$ | Voltage slope factor in GJ inactivation $n_J$ | 5 to 8 | |
| $\tau_J$ | $ms$ | Time constant of GJ inactivation $n_J$ | 100 | |
| $\rho_l$ | | Fraction of active junctions at ($i$-1)th connection | 0.8 | |
| $\rho_r$ | | Fraction of active junctions at ($i$+1)th connection | 0.2 | |
| $V_r$ | $mV$ | Physiological $V_r$ resting state | -60 to -55 | Depolarized physiological $V_r$, set to neighboring cell |
| $V_{dr}$ | $mV$ | Very depolarized $V_r$ resting state, | -25 to -10 | Very depolarized resting $V_r$, set to neighboring cell |
| $I_{ext}$ | $nA$ | External constant current | 0.5-0.6 | Bifurcation parameter |

### References

1 G. E. Sosinsky and B. J. Nicholson, "Structural organization of gap junction channels," Biochim Biophys Acta **1711** (2), 99–125 (2005).

2 J E Rash, T. Yasumura, F E. Dudek, and J. Nagy, "Cell-specific expression of connexins and evidence of restricted gap junctional coupling between glial cells and between neurons," Journal of Neuroscience **21** (6), 1983–2000 (2001).

3 S. Aten, C. M. Kiyoshi, E. P. Arzola, J. A. Patterson, A. T. Taylor, Y. Du, A. M. Guiher, M. Philip, E. G. Camacho, D. Mediratta, K. Collins, K. Boni, S. A. Garcia, R. Kumar, A. N. Drake, A. Hegazi, L. Trank, E. Benson, G. Kidd, D. Terman, and M. Zhou, "Ultrastructural view of astrocyte arborization, astrocyte-astrocyte and astrocyte-synapse contacts, intracellular vesicle-like structures, and mitochondrial network," Prog Neurobiol **213**, 102264 (2022).

4 G. Sosinsky, "Gap junction structure: New structures and new insights", in *Gap Junctions - Current topics in membranes* (Elsevier, 2000), Vol. 49, pp. 1–22.

5 R. Wilders and H. J. Jongsma, "Limitations of the dual voltage clamp method in assaying conductance and kinetics of gap junction channels," Biophys J **63** (4), 942–953 (1992).

6 H. Jongsma, R. Wilders, A.C.G. van Ginneken, and M.B. Rock, "Modulatory effect of the transcellular electrical field on gap junctional conductance", in *Biophysics of Gap Junction Channels*, edited by C. Peracchia (CRC Press, Boca Raton, FL, 1991), pp. 163–172.

7 T. A. Bargiello, S. Oh, Q. Tang, N. K. Bargiello, T. L. Dowd, and T. Kwon, "Gating of Connexin Channels by transjunctional-voltage: Conformations and models of open and closed states," Biochim Biophys Acta Biomembr **1860** (1), 22–39 (2018).

8 A. Revilla, M.V.L. Bennett, and L. C. Barrio, "Molecular determinants of membrane potential dependence in vertebrate gap junction channels," Proceedings of the National Academy of Sciences **97** (26), 14760–14765 (2000).

9 L. C. Barrio, T. Suchyna, T. Bargiello, L. X. Xu, R. S. Roginski, M. V. Bennett, and B. J. Nicholson, "Gap junctions formed by connexins 26 and 32 alone and in combination are differently affected by applied voltage," Proc Natl Acad Sci U S A **88** (19), 8410–8414 (1991).

10 P. Janjic, D. Solev, and L. Kocarev, "Non-trivial dynamics in a model of glial membrane voltage driven by open potassium pores," Biophys J **122** (8), 1470–1490 (2023).

11 M. Armbruster, S. Naskar, J. P. Garcia, M. Sommer, E. Kim, Y. Adam, P. G. Haydon, E. S. Boyden, A. E. Cohen, and C. G. Dulla, "Neuronal activity drives pathway-specific depolarization of peripheral astrocyte processes," Nat Neurosci **25** (5), 607–616 (2022).

12 E.M. Izhikevich, *Dynamical Systems in Neuroscience - The Geometry of Excitability and Bursting*. (The MIT Press, Cambridge, Massachusetts, 2007).

13 Z. Zhao and H. Gu, "Transitions between classes of neuronal excitability and bifurcations induced by autapse," Sci Rep **7** (1), 6760 (2017).

14 G. Seifert, K. Hüttmann, D.K. Binder, C. Hartmann, A. Wyczynski, C. Neusch, and C. Steinhäuser, "Analysis of Astroglial K+ Channel Expression in the Developing Hippocampus Reveals a Predominant Role of the Kir4.1 Subunit," *J. Neurosci.* **29**, 7474–7488 (2009).
